# Impaired Cx43 gap junction endocytosis causes cardiovascular defects in zebrafish

**DOI:** 10.1101/2021.03.07.434329

**Authors:** Caitlin Hyland, Michael Mfarej, Giorgos Hiotis, Sabrina Lancaster, Noelle Novak, M. Kathryn Iovine, Matthias M. Falk

**Affiliations:** Department of Biological Sciences, Lehigh University Iacocca Hall, 111 Research Drive, Bethlehem PA, 18015

**Keywords:** Cx43, gap junction, heart, zebrafish, endocytosis, vasculature, gap junction intercellular communication (GJIC)

## Abstract

Gap junction proteins, termed connexins (Cx), mediate direct cell-to-cell communication by forming channels that physically couple cells, thereby linking their cytoplasm, permitting exchange of molecules, ions, and electrical impulses. The most ubiquitously expressed gap junction protein, connexin43 (Cx43) has been implicated in cardiovascular diseases including arrhythmias, cardiomyopathies, hypertension and diabetes. The Cx43 C-terminal (CT) domain serves as the regulatory hub of the protein affecting all aspects of gap junction function. Here, deletion within the Cx43 CT (amino acids 256-289), a region known to encode key residues regulating gap junction turnover is employed to examine the effects of dysregulated Cx43 gap junction endocytosis using cultured cells (Cx43^Δ256-289^) and zebrafish model (*cx43^lh10^*). We report that this CT deletion causes defective gap junction endocytosis as well as increased gap junction intercellular communication (GJIC). Increased Cx43 protein content in cx*43^lh10^* zebrafish, specifically in the cardiac tissue, larger gap junction plaques and longer Cx43 protein half-lives coincide with severely impaired cardiovascular development. These findings suggest that normal, unimpaired Cx43 gap junction endocytosis and turnover is an essential aspect of gap junction function as demonstrated here for cardiovascular development that when impaired can give rise to arrhythmias, heart malformations and aberrant vasculature structure and function.

## INTRODUCTION

Tissues and organ systems of multicellular organisms arise through tightly regulated developmental programs coordinating cell differentiation and growth. An underpinning of developmental structuring of tissues is cell-to-cell communication. Gap junction proteins are pivotal to developmental pathways through mediation of direct intercellular communication. Gap junctions are comprised of hundreds to thousands of channels made of connexin (Cx) proteins. These channels provide a hydrophilic pore between adjacent cells which propagate chemical signals throughout tissues (Thévenin *et al*., 2013; Falk *et al*., 2014). One gap junction protein that is emerging as a critical factor in development of cardiovascular system structure and function is connexin43 (Cx43). Although Cx43 is ubiquitously expressed, Cx43 is enriched in atrial and ventricular myocytes where it localizes to cell-cell interfaces called intercalated disks (Lampe and Lau, 2004). Defects in Cx43-dependent gap junction intercellular communication (GJIC) results in severe cardiomyopathies *in vivo* including heart failure, dilated cardiomyopathy, myocardial ischemia, lethal pulmonary outflow obstruction, and lethal arrhythmias (Beardslee *et al*., 1998; Beardslee *et al*., 2000; Bruce *et al*., 2008; Duffy, 2012; Martins-Marques *et al*., 2015; Severs *et al*., 2004). These findings demonstrate that Cx43 biology represents an exciting area of research in understanding the molecular etiology of cardiomyopathies.

While the importance of Cx43 in heart development is established, and the molecular events that dictate the Cx43 life cycle have largely been characterized, the physiological role of gap junction endocytosis is not well understood. Does aberrant gap junction endocytosis cause disease? This question becomes especially important as gap junction connexins, surprisingly for a structural membrane protein have a very short, 1-5 hour half-live (Berthoud et al., 2004; Fallon and Goodenough, 1981; Chu and Doyle, 1985; Hare and Taylor, 1991), resulting in gap junction channels to constantly be endocytosed and replaced (Lauf *et al*., 2002; Gaietta *et al*., 2003; Piehl *et al*., 2007; Falk *et al*., 2009). The Cx43 C-terminus (CT), spanning residues 232-382 (232-381 in zebrafish Cx43), is a regulatory hub serving as a substrate for key post-translational modifications and accessory protein binding (Lampe *et al*., 2004; Thévenin *et al*., 2013; Falk *et al*., 2014; Thévenin *et al*., 2017; Leithe *et al*., 2018). A coordinated series of phosphorylation and dephosphorylation events on the Cx43 CT control trafficking of oligomerized connexin complexes (termed connexons or hemichannels) to the plasma membrane, recruitment and docking of connexons to gap junction plaques regulated by the scaffolding protein, zonula occludens-1 (ZO-1), and finally ubiquitin-dependent clathrin-mediated endocytosis of gap junction channels (Park *et al*., 2006; Piehl *et al*., 2007; Solan *et al*., 2007; Gumpert *et al*., 2008; Falk *et al*., 2009; Park *et al*., 2009; Rhett *et al*., 2011; Falk *et al*., 2012; Fong *et al*., 2013; Thévenin *et al*., 2013; Cone *et al*., 2014; Dunn *et al*., 2014). All of these events, when mutated, trigger complications in gap junction function, such as abundant channel accrual, aberrant channel closure, and defective gap junction endocytosis. Despite the importance of the Cx43 CT domain in regulating gap junction function, sequence analysis of mRNAs expressed in over 60,000 healthy unrelated human individuals from the Exome Aggregation Consortium database (Lek et al., 2016; Karczewski *et al*., 2020) reveal over 1,100 mutations in the Cx43 CT, however, no mutations exist at any of the key regulatory residues we and others characterized critical for gap junction endocytosis (ZO-1 binding site, D^379^LIE^382^; serine phosphorylation sites, S279/282, S365, S368, S373; K63 poly-ubiquitylation sites, K264, K303; and AP-2/clathrin binding sites S2: Y^265^AYF^268^ and S3: Y^286^KLV^290^) (Fong *et al*., 2012; Fong *et al*., 2013; Fong *et al*., 2014; Nimlamool *et al*., 2015; Thévenin *et al*., 2017; Kells-Andrews *et al*., 2018;). This finding suggests that normal, unimpaired gap junction endocytosis is essential for developmental programs, and more broadly overall gap junction function.

Here, we analyzed defects in cardiovascular development that occur due to the deletion of amino acid residues (Cx43^Δ256-289^) that are critical for Cx43-based gap junction endocytosis, including the conserved AP-2/clathrin binding site (S2), S279/282 phosphorylation sites, the binding site for the E3-ligase, Nedd4 that ubiquitinates K264 and K303, and one of the ubiquitination sites, K264 in the Cx43 CT using the zebrafish (*Danio rerio*) as a model system. Choosing zebrafish and not mice is feasible as Cx43 gap junctions have conserved roles in vertebrate embryonic development across species (Chatterjee et al., 2005), endocytic signals are largely conserved, and the transparency of zebrafish embryos readily allows the detection of developmental defects. We report that zebrafish harboring a homozygous deletion of amino acid residues 256-289 in the Cx43 CT domain (designated *cx43^lh10^*) exhibit a suite of cardiovascular phenotypes including heart malformation, bradycardia and aberrated vasculature structure and function. Cx43^Δ256-289^-dependent cardiovascular phenotypes coincide with significantly increased Cx43 protein and mRNA levels in addition to dysregulated gene expression *in vivo*. Moreover, analyses of the Cx43^Δ256-289^ mutation in a cell culture model reveal increases in gap junction plaque size, prolonged Cx43 half-life, and increased dye transfer, indicative of increased GJIC. Our findings demonstrate that undisturbed gap junction endocytosis and turnover plays a profound role for gap junction function as demonstrated here in the cardiovascular system.

## RESULTS

### *cx43^lh10^* zebrafish exhibit significantly increased Cx43 expression

To investigate the effects of impaired Cx43 gap junction endocytosis in zebrafish, we generated a zebrafish transgenic line (designated *cx43^lh10^*) in which many of the conserved key regulatory amino acid residues involved in gap junction endocytosis located in the Cx43 C-terminus are deleted (Figure. 1A) (Thévenin *et al*., 2013). These residues include the critical MAPK phosphorylation sites S261, S279, S282 (involved in channel closure and decreased GJIC), and residues triggering clathrin mediated endocytosis including K264 (ubiquitination), the Nedd4 E3-ubiquitin ligase binding site, and the conserved AP2/clathrin binding site (S2) (Falk *et al*., 2009; Girão *et al*., 2009; Fong *et al*., 2012; Fong *et al*., 2013; Thévenin *et al*., 2013; Falk *et al*., 2014; Martins-Marques *et al*., 2015; Nimlamool *et al*., 2015; Kells-Andrews *et al*., 2018) (Figure 1A, B). Note, that amino acid numbering in the zebrafish Cx43 is partially shifted one position to the left compared to Mammalia due to the deletion of an amino acid residue in the Cx43 zebrafish CT. The transgenic line was generated using the CRISPR/Cas9 gene editing system. Zebrafish embryos were microinjected at the one-cell stage of development with the CRISPR components and the efficiency of the CRISPR/Cas9 system was verified by PCR and sequencing at 24hpf, (Figure 1B, Supplemental Figure 1). The *cx43^lh10^* zebrafish, containing the *Cx43^Δ256-289^* deletion exhibits no significant decrease in viability or any evident morphological defects in early development (Supplemental Figure 2).

**FIGURE 1:**
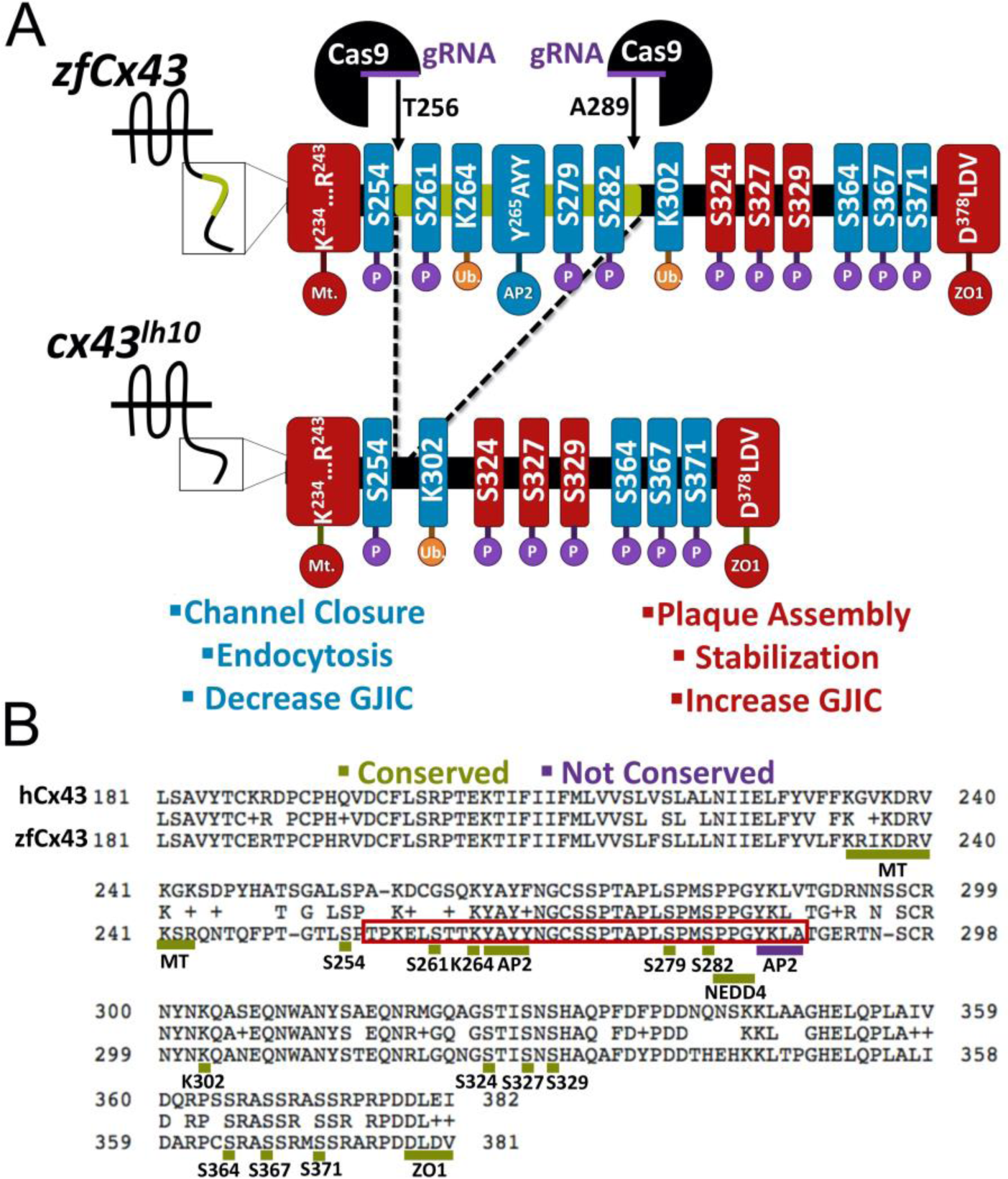
Generation of the *cx43^lh10^* zebrafish line using the CRISPR/Cas9 gene editing tool. (A) Schematic of the zebrafish WT Cx43 C-terminus (CT) (top) and *cx43^lh10^* CT (bottom). Two guide RNAs (gRNAs) were designed, one that targeted amino acid 256 and the other that targeted amino acid 289. After generation of double strand breaks, the system relies on non-homologous end joining (NHEJ) to repair the breaks, omitting amino acids 256-289 from the Cx43 gene. The portion of Cx43 deleted in *cx43^lh10^* (green) contains several key residues involved in gap junction endocytosis including the conserved AP2/clathrin binding site, critical MAPK phosphorylation sites S261 and S279/S282, and one of the two poly-ubiquitination sites (blue) whereas, residues involved in plaque assembly, stabilization, and increased GJIC (red), are left intact. (B) Amino acid sequence alignment of the CT of human Cx43 (top) and zebrafish Cx43 (bottom). The boxed region (red) encompasses the deleted amino acid residues and displays the critical endocytosis-related amino acid motifs. Underlined regions indicate conserved (green) and not conserved (purple) sequence motifs between human and zebrafish Cx43. Note, that all residues in (A) and (B) are labeled following zebrafish nomenclature, which is partially shifted one position to the left compared to mammalian Cx43 due to the deletion of one amino acid residue in the CT of zebrafish Cx43.

To assess if the deleted sequence (Cx43^Δ256-289^) in the *cx43^lh10^* zebrafish affects Cx43 gap junction endocytosis, immunofluorescence (IF) analysis of Cx43 levels was employed in *cx43^lh1^* zebrafish to assess total Cx43 protein levels (Figure 2). Whole 24hpf zebrafish were fixed and stained for Cx43 and protein levels were quantified using Image J (Schindelin *et al*., 2012). *cx43^lh10^* embryos exhibit significantly increased total Cx43 protein levels relative to wild type (WT) zebrafish embryos (Figures 2A, B), consistent with our previously reported immunoblot analyses of Cx43 protein level from regenerating fins of *cx43^lh10^* zebrafish (Bhattacharya *et al*., 2020).

**FIGURE 2:**
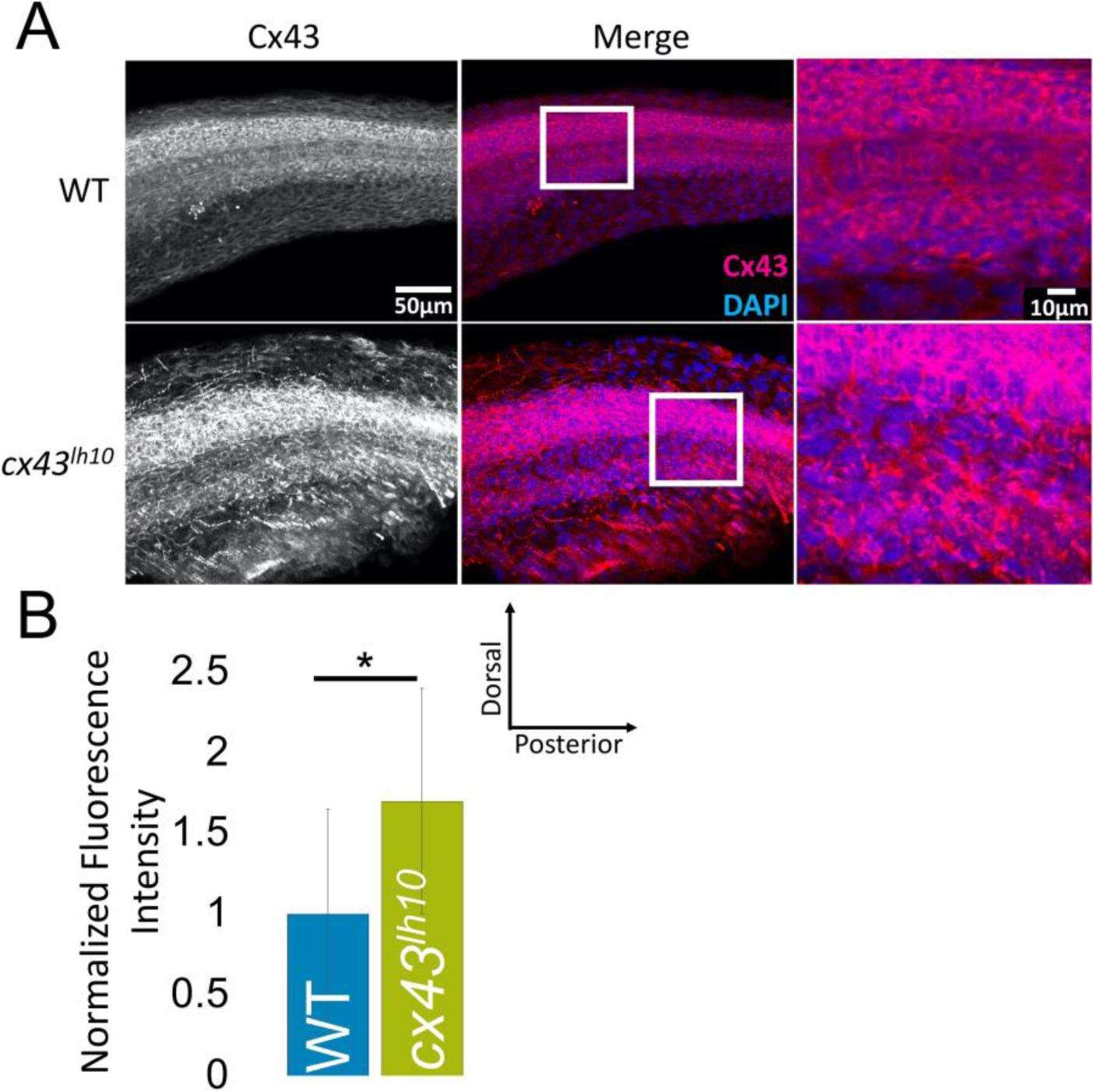
Cx43 is dysregulated in *cx43^lh10^* zebrafish. (A) Representative immunofluorescence staining of Cx43 (magenta) and DAPI (blue) in 24hpf WT (top) and *cx43^lh10^* zebrafish embryos. (B) Quantification of the immunofluorescence staining indicating increased Cx43 protein levels in *cx43^lh10^* embryos compared to WT. (n= 5 WT and *cx43^lh10^* embryos each, p=0.03). Error: s.d.

### Cx43^Δ256-289^ expressed in cultured cells exhibits significantly larger gap junction plaques, longer protein half-live, and increased dye transfer capability

The Cx43 CT residues, 256-289, contain targets which regulate Cx43 endocytosis. To assess if the observed increased protein levels in *cx43^lh10^* zebrafish are due to increased accumulation of Cx43 in the plasma membrane, constructs bearing the zebrafish coding sequence for the Cx43^Δ256-289^ allele were expressed in HeLa and MDCK cell models which do not contain endogenous Cx43. HeLa cells were transfected with either Cx43^Δ256-289^ or WT constructs and fixed 24 hours post transfection. After fixation, HeLa cells were stained for Cx43 via immunofluorescence. As expected, HeLa cells expressing Cx43^Δ256-289^ exhibited two-fold greater gap junction plaque size compared to HeLa cells transfected with WT Cx43 (Figure 3A, B). To test if increased gap junction plaque size in Cx43^Δ256-289^ expressing cells coincided with decreased Cx43 endocytosis, protein half-life analysis of Cx43^Δ256-289^ and WT Cx43 expressing HeLa cells was employed. HeLa cells were transfected with either Cx43^Δ256-289^ or WT Cx43 constructs and 24 hours post transfection, cells were treated with the ribosomal blocker, cycloheximide to halt new protein biosynthesis. HeLa cells expressing WT zebrafish Cx43 exhibit a half-life of 5-6 hours, consistent with previous findings (Fallon *et al*., 1981; Beardslee *et al*., 1998; Falk *et al*., 2009; Falk *et al*., 2014). However, HeLa cells transfected with Cx43^Δ256-289^ fail to exhibit an observable decrease in Cx43 protein levels following the 6-hour incubation period in the presence of cycloheximide (Figure 3C and 3D), indicating a significantly longer half-life of the Cx43^Δ256-289^ protein. This is consistent with a similar mutant (Δ254-290) that we constructed and tested previously (Fong *et al*., 2013). To test whether Cx43^Δ256-289^ expressing cells have altered GJIC, we performed scrape-loading Lucifer Yellow (LY) dye transfer assays in Cx43^Δ256-289^ and WT Cx43 expressing MDCK cells. The distance of LY (a gap junction permeable dye) transferring between cells, through gap junctions, away from the site of injury (scrape) was determined by quantitatively measuring LY-fluorescence signals. LY transfer in cells expressing WT Cx43 is significantly increased compared to the untransfected control (no gap junctions). In addition, Cx43^Δ256-289^ expressing MDCK cells exhibited significantly further increased dye transfer compared to MDCK cells expressing WT Cx43 (Figure 3E, F). This indicates that cells expressing Cx43^Δ256-289^ are capable of generating functional gap junction channels with increased GJIC. This result is consistent with our previous findings suggesting increased GJIC in regenerating fins in *cx43^lh10^* zebrafish (Bhattacharya *et al*., 2020). Taken together, the increased half-life, and the observed increase in the size of gap junction plaques, combined with increased dye transfer in cells expressing Cx43^Δ256-289^ suggests aberrant Cx43 gap junction endocytosis, as well as increased GJIC in *cx43^lh10^* zebrafish and is consistent with a role for residues 256-289 in the regulation of Cx43 endocytosis (Fong *et al*., 2013; Thévenin *et al*., 2017).

**FIGURE 3:**
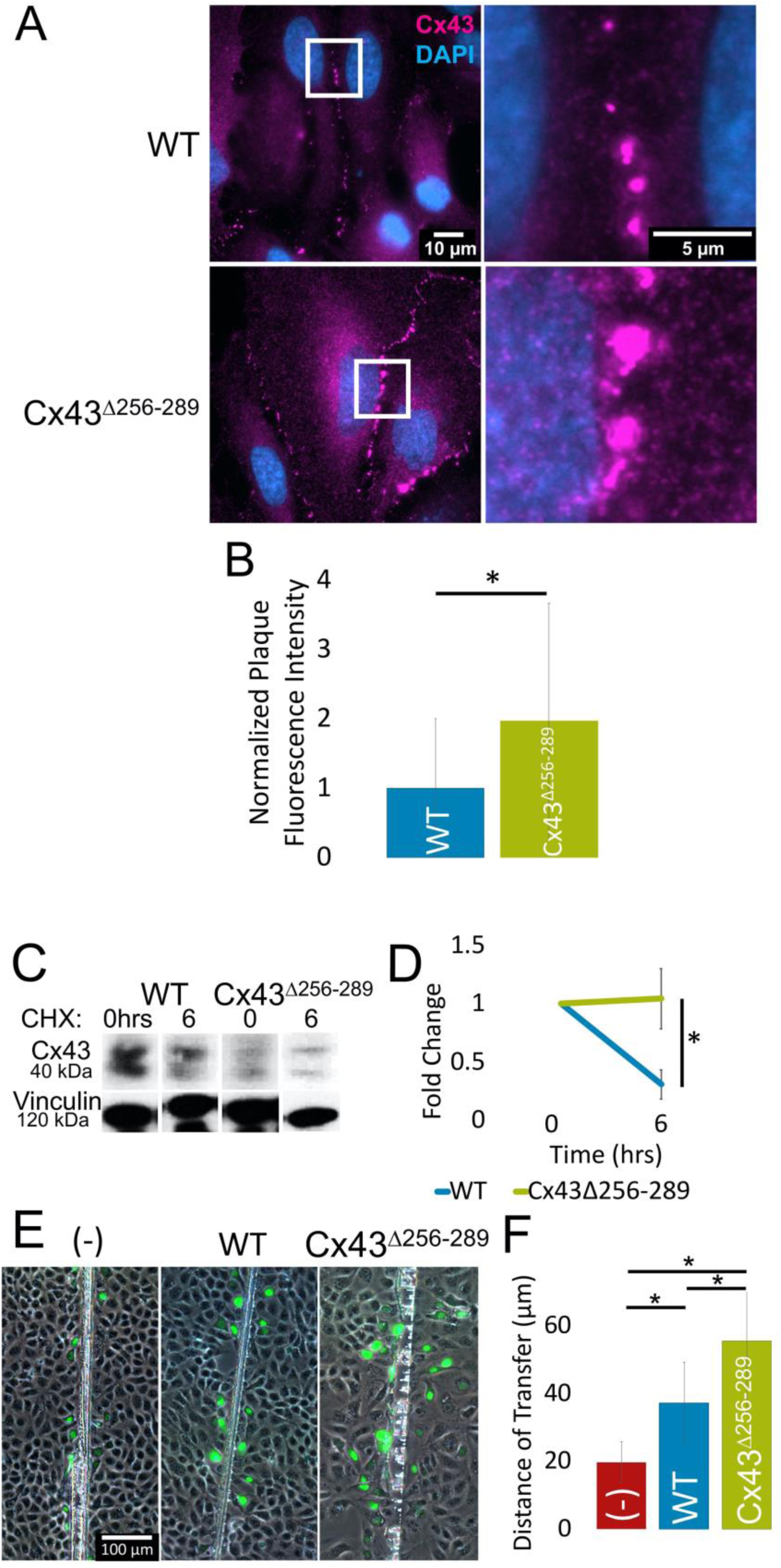
zfCx43^Δ256-289^ exhibits larger plaques, longer half-lives, and increased dye transfer in cultured cells. (A, B) Representative immunofluorescence and quantitative plaque size analysis of Cx43 (magenta) and DAPI (blue) showing gap junction plaques in the membranes of HeLa cells. Plaque size analysis reveals that Cx43^Δ256-289^ expressing cells exhibit larger gap junction plaques compared to WT Cx43 expressing cells. n=258 WT and 190 Cx43^Δ256-289^ plaques, p=2.1x10^-5^. (C, D) Representative western blot of Cx43 protein half-life at 0hrs and 6hrs after cycloheximide treatment and quantitative analysis. Western blot quantification indicates Cx43^Δ256-289^ protein is significantly less degraded 6 hours post cycloheximide treatment than WT Cx43 protein. n=3 transfected cultures, p=0.02. (E, F) Representative images of scrape-loading dye transfer assay of untransfected MDCK cells (-) (red), MDCK cells transfected with WT Cx43 (blue), and MDCK cells transfected with Cx43^Δ256-289^ (green). Quantification indicates that cells expressing Cx43^Δ256-289^ exhibit significantly increased dye transfer compared to cells expressing WT Cx43 or cells with no Cx43 protein. n= 3 transfected cultures, p=0.006 (WT v. (-)), 2.4x10^-10^ (WT v. Cx43^Δ256-289^), 7.4x10^-18^ ((-) v. Cx43^Δ256-289^). Error: s.d.

### Increased Cx43 protein in cardiac tissue of *cx43^lh10^* zebrafish results in larger gap junction plaques

Given the role of Cx43 in cardiac function and the growing evidence suggesting the CT is essential for cardiac function (Beardslee *et al*., 1998; Maass *et al*., 2007; Bruce *et al*., 2008; Maass *et al*., 2009; Chi *et al*., 2010; Duffy, 2012; Lübkemeier *et al*., 2013; Martins-Marques *et al*., 2015a; Martins-Marques *et al*., 2015b; Martins-Marques *et al*., 2015c; Martins-Marques *et al*., 2015d), it became important to test the role of the *cx43^lh10^* mutation in cardiovascular development and physiology in *cx43^lh10^* zebrafish, as zebrafish offers insight that is inaccessible in mice due to the ability to readily observe cardiac function in live embryos.

Immunofluorescence analysis of dissected cardiac tissue from *cx43^lh10^* zebrafish revealed a ∼1.3-fold and ∼1.4-fold increase in Cx43 levels in the *cx43^lh10^* zebrafish ventricle and atria, respectively, relative to WT zebrafish (Figure 4A, B), which is consistent with increased Cx43 in developing *cx43^lh10^* zebrafish embryos (Figure 2A, B) and in adult *cx43^lh10^* zebrafish caudal fins (Bhattacharya *et al*. 2020). Moreover, increased Cx43 levels in *cx43^lh10^* zebrafish hearts coincided with ∼3.4-fold and ∼2.7-fold increase in gap junction plaque size in the cx*43^lh10^* zebrafish ventricle and atria, respectively, relative to WT zebrafish (Figure 4A, C), indicating decreased gap junction endocytosis in cardiac tissue of *cx43^lh10^* zebrafish, likely causing increased GJIC, as seen in the caudal fin (Bhattacharya *et al*., 2020) and in Cx43^Δ256-289^ expressing MDCK cells.

**FIGURE 4:**
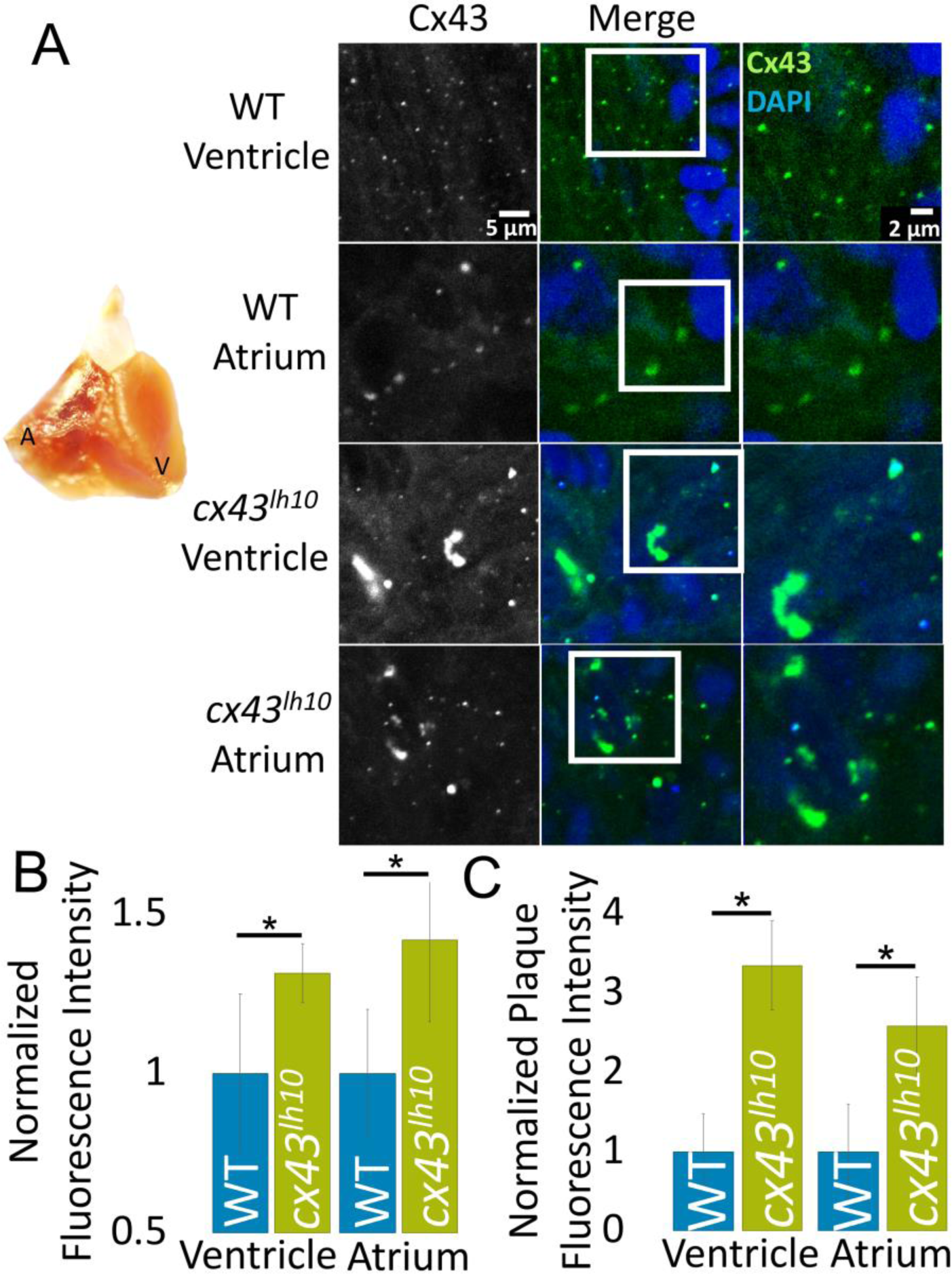
Cx43 protein and gap junction size is increased in *cx43^lh10^* zebrafish embryos. (A) Representative immunofluorescence of Cx43 (green) and DAPI (blue) in the atrium and ventricle of dissected WT and *cx43^lh10^* adult zebrafish hearts. Dissected WT zebrafish heart (left) indicating location of atrium and ventricle (also see Figure 5A). (B) Cx43 protein level quantification. n=3 WT and *cx43^lh10^* hearts, p=0.005. (C) Plaque size analysis of WT compared to *cx43^lh10^* cardiac gap junctions. n=3 WT and *cx43^lh10^* hearts, p=1.77x10^-5^. Error: s.e.m.

### *cx43^lh10^* zebrafish have altered cardiac function and morphology

Altered Cx43 function and GJIC is associated with several cardiomyopathies including heart failure, myocardial ischemia, and cardiac arrythmias (Beardslee *et al*., 2000; Bruce *et al*., 2008; Duffy, 2012; Martins-Marques *et al*., 2015d). If *cx43^lh10^* zebrafish have impaired gap junction endocytosis and GJIC in cardiac tissue, then increased atrial and ventricular Cx43 levels should result in altered cardiac morphology and/or function. To test this, heart rates were measured in live 3dpf and 6dpf embryos. WT embryos exhibit an average heart rate of about 155bpm at 3dpf and 180bpm at 6dpf, whereas *cx43^lh10^* embryos have an average heart rate of about 145bpm at 3dpf and 165bpm at 6dpf (Figure 5B, Movies 1 and 2). In addition to decreased heart rate, a significantly larger portion of *cx43^lh10^* embryos develop elongated, malformed hearts and pericardial edema (Figures 5A, C). This data suggests that reduced Cx43 endocytosis and associated increased GJIC leads to abnormal cardiovascular development and bradycardia.

**FIGURE 5:**
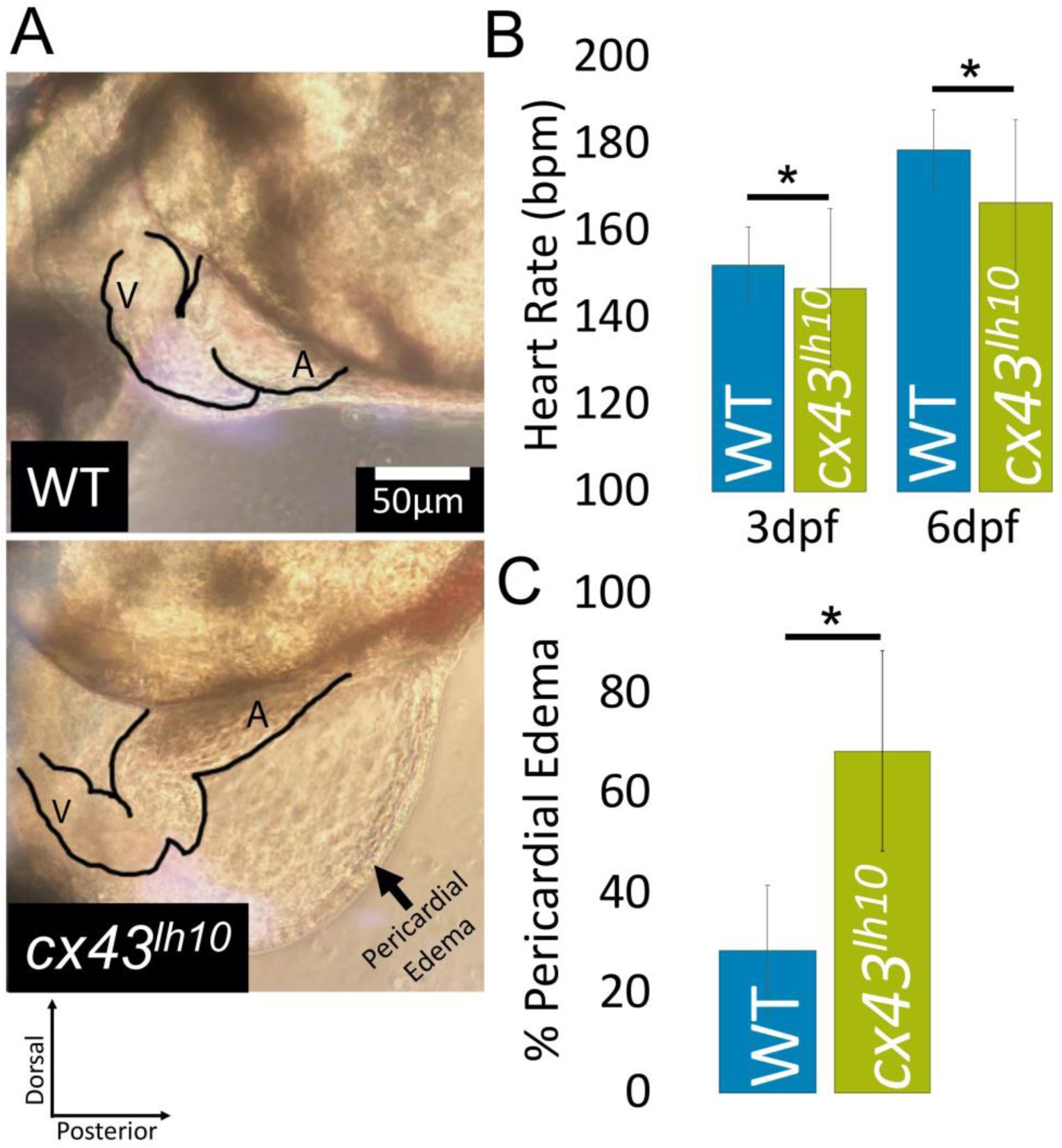
*cx43^lh10^* zebrafish exhibit cardiac defects. (A) Cardiac morphology of WT vs *cx43^lh10^* 3dpf zebrafish. (B) Heart rates of WT and *cx43^lh10^* zebrafish at 3dpf and 6dpf. n= 100 WT and *cx43^lh10^* embryos (3dpf, 6dpf) each, p=0.0009 (3dpf), 10x10^-13^ (6dpf). (C) Percentage of embryos exhibiting pericardial edema at 3dpf. n=496 (WT) and 468 (*cx43^lh10^*) 3dpf embryos, p=0.04. Error: s.d.

### *cx43^lh10^* zebrafish have malformed and disorganized vasculature

Given the cardiac phenotypes exhibited in *cx43^lh10^* fish, we thought to determine if the *cx43^lh10^* fish also exhibit vascular abnormalities. Cx43 is primarily expressed in vasculature smooth muscle cells and the endothelial cells of the vasculature. Cx43 coordinates synchronous vasomotor tone, vessel constriction and cell proliferation and migration in the vasculature (Figueroa *et al*., 2008). To determine if there are any vasculature abnormalities in *cx43^lh10^* zebrafish, we investigated the intersegmental vessels in developing embryos (5dpf) by cross-breeding our *cx43^lh10^* line with the TG(fli1:EGFP) transgenic line that drives GFP expression in all vasculature throughout development (Lawson *et al*., 2002). We found that the organization of the developing vasculature in *cx43^lh10^* embryos appears largely normal, however vessel diameter is significantly larger in diameter than in WT vasculature (Figure 6A, B).

**FIGURE 6:**
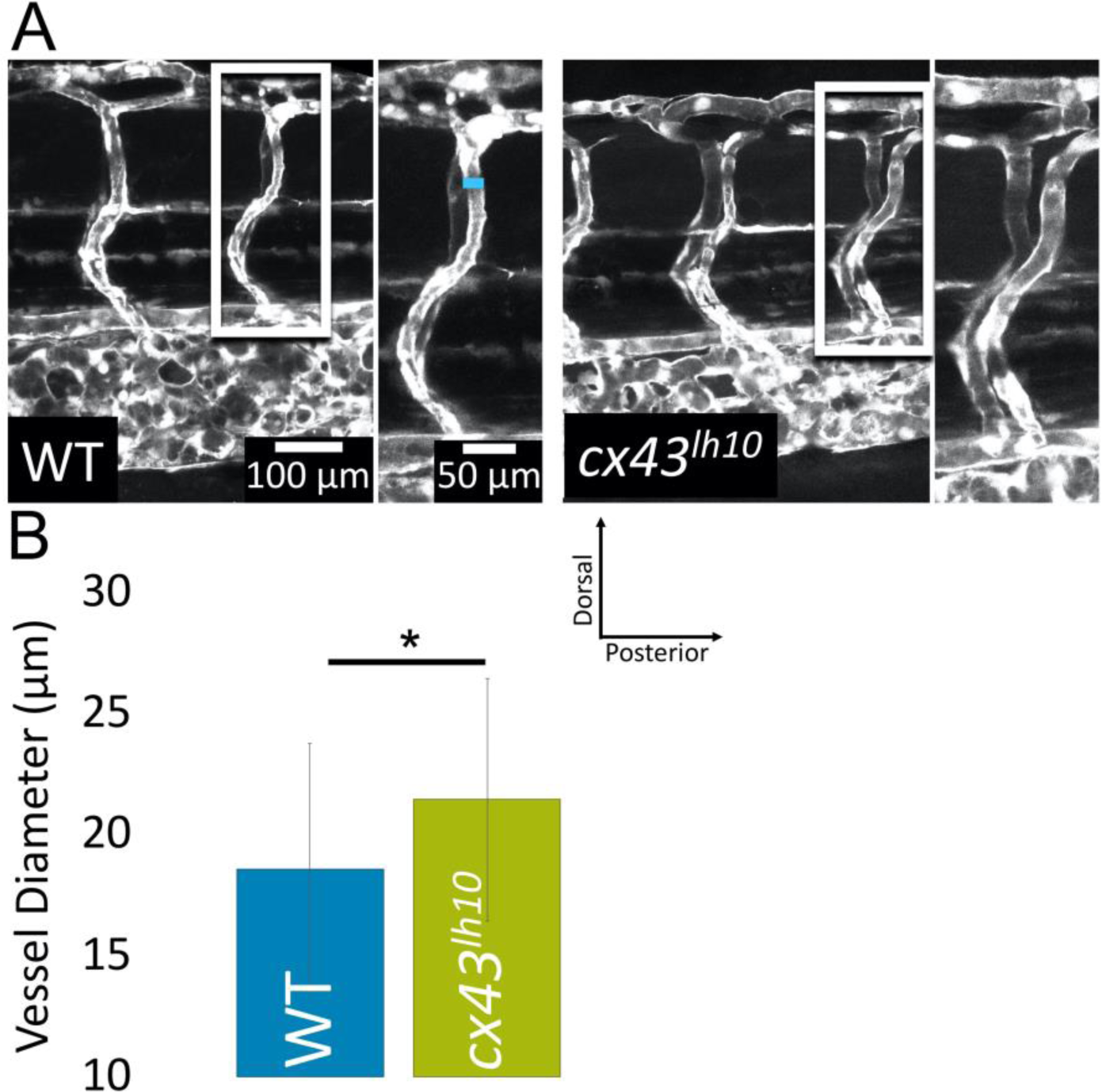
*cx43^lh10^* zebrafish embryos exhibit defects of the vasculature. (A) GFP-tagged vasculature of 5dpf TG(fli1:EGFP) (left) and *cx43^lh10^*/TG(fli1:EGFP) (right) embryos, emphasizing intersegmental vessels. Diameter measurement indicated by blue line in (A). (B) Quantification of vessel diameter in 5dpf embryo vasculature. n= 5 (WT), 3 (lh10), p=0.0003. Error: s.d.

To further test for vascular organization and function, we analyzed the structure, organization and function of the vasculature of adult *cx43^lh10^* zebrafish in caudal fins. Anesthetized adult fish were used for blood vessel diameter measurements and blood flow recordings in the vasculature. In WT adult fins, veins are organized flanking each fin ray bone and an artery extends the middle of the ray bone between the two veins (Figure 7A). In WT zebrafish, veins are typically 2 times the diameter of arteries (Figure 7A, B) and blood flow is significantly slower in veins than in arteries (Figure 7E, Movie 3). In contrast, *cx43^lh10^* arteries are significantly larger than arteries in WT fins, and veins are significantly smaller than WT veins. Moreover, while WT veins are larger than WT arteries, *cx43^lh10^* veins and arteries are of similar size (Figure 7B). In addition, arteries and veins are profoundly disorganized in the adult fin of *cx43^lh10^* fish. Based on the direction of blood flow, in many cases, arteries are positioned flanking fin rays, where normally veins are positioned in WT fish. Also, veins are positioned along the middle of the fin ray, where arteries are normally positioned in WT fish (Figure 7C, Movies 3, 4). Moreover, vasculature malformation in *cx43^lh10^* fish results in blood vessel dysfunction as determined by drastically reduced blood flow in many blood vessels in *cx43^lh10^* adult fins to the point where, in some cases, blood flow within vessels was completely halted (Figure 7D, 7E, Movies 4, 5). This vascular malformation may represent as a secondary effect caused by the described heart function defects. However, since proper gap junction coupling is required for maintaining vasomotor tone, these vasculature phenotypes in *cx43^lh10^* fish may be the result of aberrant GJIC between the cells of the vasculature due to dysregulated Cx43 endocytosis. These results could therefore support a role for Cx43 turnover in vasculature structuring and function.

**FIGURE 7:**
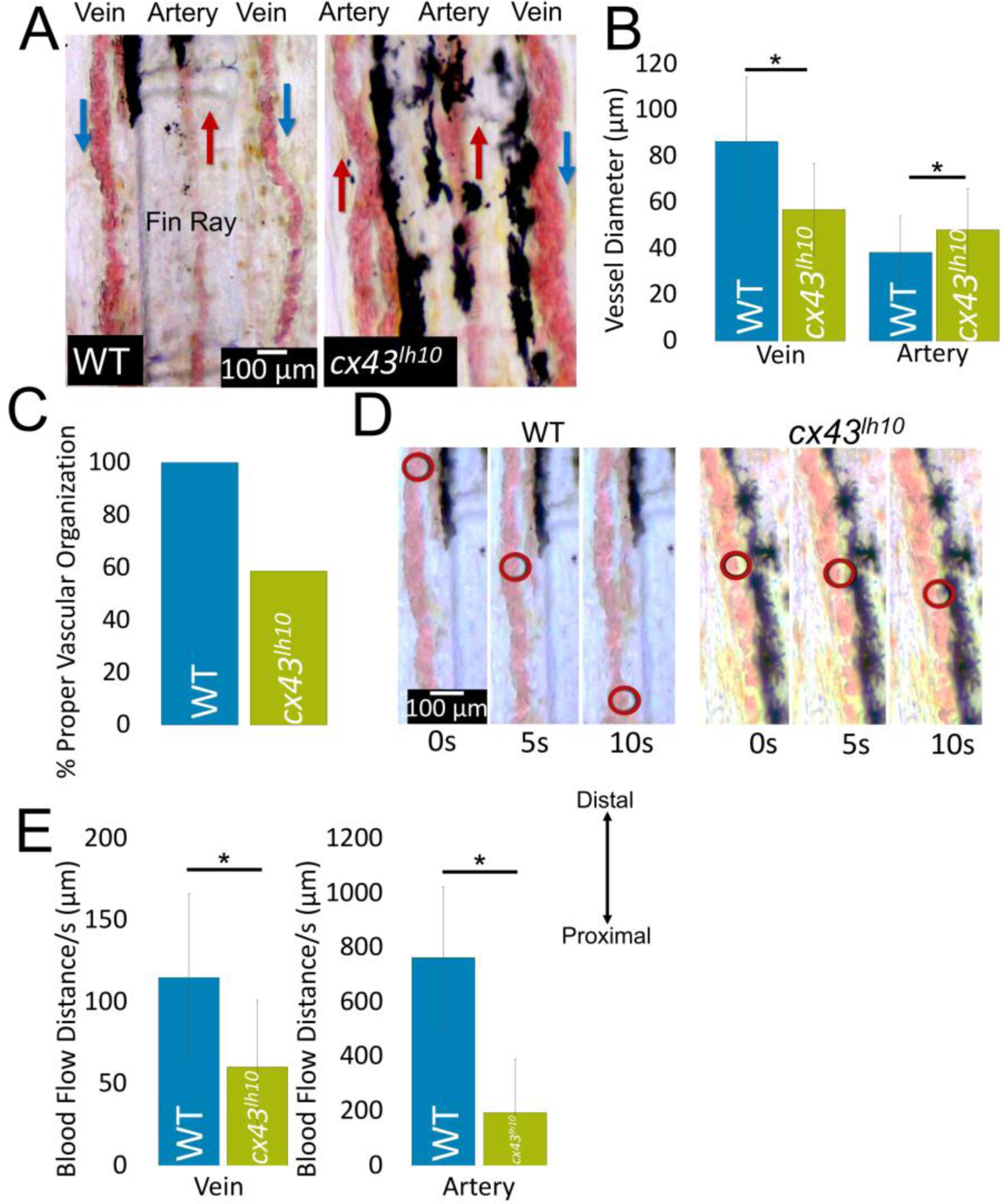
Adult *cx43^lh10^* zebrafish exhibit defective and unorganized vasculature in caudal fins. (A) WT and *cx43^lh10^* vasculature in the adult fin, highlighting the typical organization of the vasculature in the fin (WT) and the vascular disorganization seen in *cx43^lh10^* fish. (B) Artery vs vein diameter in WT fins and *cx43^lh10^* adult fins. n= 14 fish (WT), 17 (*cx43^lh10^*), p=0.008 (vein), 0.002 (artery). (C) % proper vascular localization in WT (100%) and *cx43^lh10^* (59%). Vasculature were characterized as either veins or arteries dependent on blood flow direction. n=14 WT fish, 17 (*cx43^lh10^*). (WT) (D) Distance of blood traveled through veins over a 10s period in WT and *cx43^lh10^* fins. (E) Average blood flow distance/s in WT and cx43*^lh10^* veins and arteries. n= 10 (WT, *cx43^lh10^*), p=0.009 (vein), 6x10^-6^ (artery). Error: s.d.

### *cx43^lh10^* fish exhibit dysregulated gene expression

In addition to functional defects in the heart and vasculature, the *cx43^lh10^* mutation may promote compensatory molecular responses in zebrafish cells expressing the *cx43^lh10^* allele to cope with the physiological stresses of this deletion. As, based on our data the *cx43^lh10^* mutation results in decreased Cx43 endocytosis and ensuing dysregulated GJIC, cells may respond through transcriptional repression of Cx43 mRNA levels to recover GJIC regulation. To test this hypothesis, qRT-PCR was employed to analyze Cx43 mRNA levels in *cx43^lh10^* zebrafish relative to WT zebrafish. Surprisingly, we found that *cx43* mRNA in the *cx43^lh10^* line is ∼1.5-fold increased relative to WT zebrafish suggesting that transcriptional repression of *cx43* mRNA is not involved in the cellular response to the Cx43^Δ256-289^ allele (Figure 8). Earlier work in mice tumor models demonstrated that Cx43 mRNA levels are inversely related to mRNA expression of the angiogenesis signaling molecule Vascular Endothelial Growth Factor (VEGF) (Suarez *et al*., 2001; Nimlamool *et al*., 2015). Given abnormal blood vessel structuring in *cx43^lh10^* zebrafish, it became important to test expression changes in *vegf* mRNA in response to the Cx43^Δ256-289^ allele.

**FIGURE 8:**
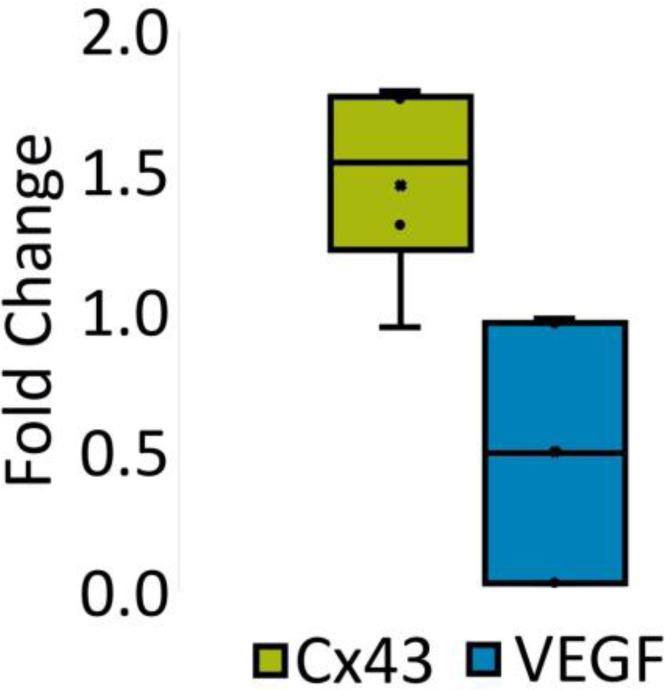
cx43*^lh10^* fish exhibit dysregulated vegf gene expression. Box and whisker plot depicting qRT-PCR fold changes of Cx43 (green) and VEGF (blue) where the “X” for each box and whisker plot indicates the average fold change for each gene (n=5, average fold changes: Cx43: 1.4, VEGF: 0.5).

Interestingly, *vegf* mRNA levels were found to be decreased ∼0.5-fold in *cx43^lh10^* zebrafish relative to WT zebrafish which is consistent with earlier findings in mice tumor models that decreased Cx43 mRNA -is associated with upregulated *vegf* mRNA expression (McLachlan *et al*., 2006; Wang *et al*., 2014) (Figure 8). These results may point to compensatory transcriptional changes in response to the *cx43^lh10^* mutation to achieve repressed VEGF expression in order to counteract arterial hypertrophy and venous hypotrophy seen in *cx43^lh10^* zebrafish (Figure 7).

## DISCUSSION

Gap junction mediated cell-to-cell communication is crucial for multicellular organisms, and when gap junctions become dysregulated a variety of damaging consequences may be encountered. The Cx43 CT harbors key regulatory residues that control gap junction endocytosis (Thévenin *et al*. 2013). We previously reported that mutating/removal of specific tyrosine-based sorting signals and other key endocytic residues located in the Cx43 CT (amino acids 254-290) abolishes clathrin association with Cx43 gap junctions, resulting in larger gap junction plaques with decreased endocytic kinetics and longer Cx43 half-life (Fong *et al*., 2013). In addition, in a recent study, we found that *cx43^lh10^* zebrafish (deletion of amino acids 256-289) exhibit increased Cx43 protein, as well as increased fin regenerate and segment length compared to WT fins. It was also found that with the addition of Gap27, a gap junction channel blocker, the phenotype was rescued, suggesting *cx43^lh10^* fish exhibit increased GJIC compared to WT (Bhattacharya *et al*., 2020). The first major finding of this study is that the *cx43^lh10^* mutant investigated here exhibits an increase in Cx43 protein levels in zebrafish which coincides with larger plaque size, a longer Cx43 half-life, and increased dye transfer in cultured cells. Taken together, this major finding indicates that removing amino acids 256-289, similarly to previous reports, causes impaired gap junction endocytosis, and an increase in GJIC.

The second major finding of this study is that *cx43^lh10^* zebrafish, besides other phenotypes exhibit severe defects of the cardiovascular system including malformed, elongated hearts, decreased heart rate, disorganized and malformed vasculature, and impaired blood flow, demonstrating that undisturbed gap junction endocytosis is crucial for normal organ development. Previous findings have shown that deletion of Cx43 in mice causes defects in the interface between the right ventricle and outflow tract, which results in blood flow obstruction and ultimately neonatal lethality due to delayed cardiac looping and malformation (Reaume *et al*., 1995; Ya *et al*., 1998). A premature stop codon, preventing the translation of most of the Cx43 CT (K258Stop) has been characterized to generate functional channels, with larger plaques at the intercalated disc, and no abnormalities in heart morphology, yet increased infarct size and susceptibility to arrhythmias were reported (Maass *et al*., 2007; Maass *et al*., 2009).

Furthermore, mice lacking the last five CT amino acid residues (Cx43Δ378stop), abolishing ZO-1 binding, displayed aberrant cardiac electrical activation properties that ultimately lead to lethal ventricular arrythmias, yet channel function was not impaired, detailing Cx43-dependent cardiac defects that are independent of channel function (Lübkemeier *et al*., 2013). It has also been shown that changes in Cx43 expression, phosphorylation, and distribution leads to cardiac arrhythmias, such as atrial fibrillation, and diseases of the myocardium such as hypertrophic cardiomyopathy and ischemic cardiomyopathy (Michela *et al*., 2015). Furthermore, defective Cx43 plays a role in diseases of the vasculature, such as hypertension and diabetes (Inoguchi *et al*., 2001; Rummery *et al*., 2004; Zhang *et al*., 2005; Figueroa *et al*., 2006; Figueroa *et al*., 2008). However, there has been very little evidence to support the mechanism/s behind defective gap junction function that leads to these disorders. *cx43^lh10^* zebrafish exhibit bradycardia, malformed hearts and pericardial edema. Pericardial edema suggests cardiac insufficiency, whereas the malformed heart might indicate an issue with cell migration, leading to cardiac looping and chamber ballooning. Once the heart tube is formed during development, the tube undergoes looping, in which the ventricle and the atrium become distinguishable with inner and outer curvatures. Then, the heart undergoes chamber ballooning, which allows for each chamber to expand (Bakkers, 2011). Previous work has shown that mechanical and electrical forces are required for proper chamber formation. For example, in zebrafish with deficiencies in ventricular or atrial contraction, the resultant phenotype includes larger, elongated chambers (Chi *et al*., 2010), similar to what is seen in the *cx43^lh10^* zebrafish. *cx43^lh10^* fish also exhibit severe defects of the vasculature including impaired blood flow, as well as disorganized and malformed vasculature. It has been previously reported that increased Cx43 expression and GJIC signals endothelial cells of the vasculature to migrate due to the necessity of establishing a continuous covering during formation and renewal of blood vessels and wound repair (Pepper *et al*., 1992a; Pepper *et al*., 1992b; Kwak *et al*., 2001; Haefliger *et al*., 2004). In addition, hypertension mouse models reveal increased Cx43 in the membranes of vascular smooth muscle cells and cardiovascular hypertrophy (Haefliger *et al*., 2004). To this end, the data presented here suggests that defective gap junction endocytosis should be considered a possible mechanism for many of these common phenotypes that are associated with mutated or dysregulated Cx43 in the cardiovascular system.

While the data presented here provides evidence for Cx43 regulatory reactions in cardiovascular development, our findings also reveal a complex interplay between Cx43, and other molecules implicated in GJIC. For instance, we report that mutations resulting in dysregulation of Cx43 endocytosis promotes upregulated *cx43* transcription which coincides with repressed transcription of *vegf*. This data suggests that VEGF may be an important player in cardiovascular development functioning downstream of Cx43 and implicates this regulatory activity in the disrupted vasculature seen in the *cx43^lh10^* mutant zebrafish. However, the putative role of altered *vegf* transcription in arterial hypertrophy and venous hypotrophy due to the *cx43^lh10^* mutation remains to be explored. One possibility is that dysregulated Cx43 endocytosis directly leads to vasculature malformations which induces *vegf* transcription modulation to compensate for aberrant vasculature patterning. This mechanism may be initiated in response to increased artery growth but also affects structuring of veins leading to drastic reductions in venous lumen size. Another possibility is that Cx43-dependent GJIC directly influences transcriptional regulators of *vegf*, but this response bears different consequences in arteries compared to veins. Separately, another interesting revelation of this study is that *cx43^lh10^* early embryos (24hpf) exhibit unaltered overall morphology, and *cx43^lh10^* fish develop into adulthood with no variations in viability despite profound cardiovascular disorders. Cx43 is expressed as early as 1.25hrs (8-cell stage) post-fertilization (Hardy *et al*., 1996; Chatterjee *et al*., 2005; Cofre *et al*., 2007) but morphological abnormalities are only apparent beginning at 24hrs post-fertilization in *cx43^lh10^* embryos. One possible explanation for this delayed onset is that other Cxs could be compensating for aberrant Cx43 function. For example, several Cx43 knockout phenotypes in mice can be rescued through replacement with other Cx types (Reaume *et al*., 1995; Plum *et al*., 2000; Gutstein *et al*., 2001; Brehm *et al*., 2007; Sridharan *et al*., 2007; Winterhager *et al*., 2007; Bedner *et al*., 2012). Mechanistic insight into the cellular response to dysregulated Cx43 endocytosis and the molecules involved in this response will generate deeper understanding for the role of regulated Cx43 endocytosis in organ system development. Taken together, the data presented here highlight the importance of undisturbed Cx43 gap junction endocytosis as a critical factor of normal gap junction function that should be considered as a possible mechanism for causing gap junction related disease.

## MATERIALS AND METHODS

### Fish maintenance

Zebrafish (*Danio rerio*) were maintained in the water system built by Aquatic Habitats (now Pentair) at 27°C -28°C temperature in a 14:10 light: dark period (Westerfield, 2007). Water quality was monitored and dosed to maintain conductivity (400–600 ms) and pH (6.95– 8/27/2019 approval). Research was performed per the IACUC for Lehigh University (protocol #231). Food was provided three times a day. Brine shrimp (hatched from INVE artemia cysts) was fed once and flake food (Aquatox AX5) supplemented with 7.5% micropellets (Hikari), 7.5% Golden Pearl (300–500 micron, Brine Shrimp direct), and 5% Cyclo-Peeze (Argent) twice per day.

### Generation of *cx43^lh10^* transgenic fish

A transgenic zebrafish line was generated that deletes amino acids 256-289 in the CT of Cx43 using the CRISPR/Cas9 genome editing tool. To generate the deletion, 2 gRNAs were designed using the ChopChop web tool (Montague *et al*., 2014; Labun *et al*., 2016; Labun *et al*., 2019). One gRNA was designed to target the C-terminal T256 region and the other gRNA was designed to target the C-terminal A289 region of the gene. Designed gRNAs were purchased from Genscript’s synthetic gRNA and crRNA service (gRNA T256: UAGACAGUUCCUUCGGCGUG; gRNA A289: CCAGGCUACAAACUGGCCAC) (Cat. No. SC1838, SC1933). Cas9 protein was purchased from Genscript (Cat. No. Z03389). Cas9 protein (12 µg) and each gRNA (30 pmol) were co-injected into one-cell-stage wild type (WT) zebrafish embryos. At 24hpf, 20% of embryos were genotyped using PCR. The remaining embryos were raised to adulthood and outcrossed to WT fish for detection of germline transmission. Heterozygous embryos were then raised to adulthood and intercrossed to generate homozygous Δ256-289 mutant zebrafish (designated *cx43^lh10^*) and the progeny sequenced.

### Generation of zebrafish wild type Cx43 and Cx43^Δ256-289^ mutant mammalian cell culture expression plasmids

Untagged zebrafish WT Cx43 and Cx43^Δ256-289^ plasmids were generated using the pEGFP-N1 vector including the full-length rat Cx43 cDNA (described in Fong *et al*., 2013) as the template. The entire rat Cx43 sequence was removed using EcoRI (Cat. No. R3101S; NEB) and BamHI (Cat. No. R3136S; NEB) restriction enzymes. The linearized and empty pEGFP-N1 vector was extracted and purified using Qiagen’s QIAquick gel extraction kit (Cat. No. 28704) for use as the backbone for the constructs. Gibson assembly (Cat. No. E5510S; NEB) was used to replace the full-length rat Cx43 in the pEGFP-N1 vector with either full-length zebrafish Cx43 obtained via PCR amplification from WT zebrafish, or Cx43^Δ256-289^ obtained via PCR amplification from *cx43^lh10^* zebrafish. Constructs were transformed into DH5α competent *E. coli* cells. Plasmids were purified using a Midiprep plasmid DNA purification kit (Cat. No. 12143; Qiagen). All constructs were verified via DNA sequence analyses.

### Whole-mount immunofluorescence of zebrafish embryos/hearts and confocal microscopy

Zebrafish embryos were fixed at 24hpf using 4% paraformaldehyde overnight at 4°C. Adult zebrafish were euthanized in an excess of 0.1% Tricaine solution and fixed in 4% paraformaldehyde overnight and hearts dissected according to Singleman *et al*. (2011). To improve staining, antigen retrieval was implemented using 1% SDS/PBS for 5 minutes while rocking at room temperature immediately before immunostaining. Embryos/hearts were washed in PBST (PBS + 0.1% Tween) and, for 1 hour prior to blocking, PBSTX (PBS + 0.1% Tween + 0.1% Triton X-100). Embryos/hearts were blocked for 1 hour using 10% BSA (Cat. No. A7906; Sigma)/PBSTX and incubated for 48 hours at 4°C in rabbit anti-Cx43 primary antibody (Cat. No. 3512; Cell Signaling) diluted 1:100 in 1% BSA/PBSTX. Embryos/hearts were then washed with 1% BSA/PBSTX and incubated for 48 hours at 4°C in goat anti-rabbit Alexa Fluor 568 (embryos) (Cat. No. A-11036; Invitrogen) or goat anti-rabbit Alexa Fluor 488 (hearts) (Cat. No. A-32731; Invitrogen) diluted 1:100 in 1% BSA/PBSTX. Embryos/hearts were also stained with 1µg/mg DAPI (Cat. No. D1306; Molecular Probes). Embryos/hearts were washed with PBST and mounted onto glass slides with 3% methylcellulose and z-stacks were generated with confocal microscopy (Zeiss LSM 880) at 25x (embryos) or 63X (hearts) using Argon lasers 568, 488, and 405nms. To quantify Cx43 protein levels, fluorescence intensity/outlined region was measured in 50 separate regions in each image of the z-stack via ImageJ (National Institutes of Health, Bethesda, MD) using the rectangle tool and averaged. Five biological replicates were performed in embryos and three biological replicates were performed in hearts.

### Cell culture and transient transfections

HeLa (gap junction deficient; Cat. No. CCL2; ATCC) and Madine-Darby Canine Kidney (MDCK) cells (gap junction deficient, Cat. No. NBL-2; ATCC) were maintained in low glucose Dulbecco’s Modified Eagle Medium (DMEM) (Cat. No. SH30021.01; Hyclone) supplemented with 50 I.U/mL penicillin and 50µg/ml streptomycin (Cat. No. 30-001-C1; Corning), 2mM L-glutamine (Cat. No. 25-005-C1; Mediatech), and 10% Fetal Bovine Serum (FBS) (Cat. No. S11150; Atlanta Biologicals) at 37°C, 5% CO_2_, and 100% humidity. Cells were washed with 1X PBS and treated with 0.25% trypsin/0.1% EDTA (Cat. No. 25-053-CI; Corning) for passaging. 24-48 hours after passaging, 60-80% confluent HeLa cells were transiently transfected with WT Cx43 or Cx43^Δ256-289^ constructs using Lipofectamine2000 (Cat. No. 11668019; Invitrogen) according to manufacturer’s recommendations.

### Cx43 plaque size analyses in HeLa cells

Cx43 gap junction plaques in HeLa cells (Cat. No. CCL2; ATCC) were analyzed via immunofluorescence and plaque size quantification. HeLa cells were grown on pretreated poly-L-lysine (Cat. No. P8920; Sigma) coated coverslips in low glucose DMEM at 37°C, 5% CO_2_, and 100% humidity. Cells were transiently transfected with either WT Cx43 or Cx43^Δ256-289^ constructs. Cells were fixed and permeabilized in ice-cold methanol. Cells were blocked in 10% FBS/PBS for 30 minutes at room temperature and incubated with primary rabbit anti-Cx43 antibodies diluted 1:200 in 10% FBS/PBS and incubated overnight at 4°C. Cells were incubated in goat anti-rabbit Alexa Fluor 568 diluted 1:200 in 10% FBS/PBS and incubated for 1 hour at room temperature. Cells were also stained with 1µg/mg DAPI. Coverslips were mounted using Fluoromount G (Cat. No. 0100-01; Southern Biotechnology) and imaged using a 60X objective on a Nikon Eclipse TE2000 inverted fluorescence microscope. To quantify gap junction plaque size, all plaques (258 in WT expressing cells and 190 in Cx43^Δ256-289^ expressing cells) in 10 cell pairs/transient transfection were outlined using the freeform tool in ImageJ and fluorescence intensity/outlined area was quantified for cell pairs expressing Cx43. Three biological replicates were performed of each construct.

### Cx43 protein half-life analyses and immunoblotting

24 hours post transfection with either the WT Cx43 or Cx43^Δ256-289^ constructs, HeLa cells were treated with 50µg/mL cycloheximide (Cat. No. C655; Sigma) for up to 6 hours at 37°C, 5% O2, and 100% humidity. Cells were lysed in SDS-PAGE sample buffer at hour 0 and hour 6 and boiled for 5 minutes. Samples were separated on 12% SDS-PAGE mini gels (BioRad). Proteins were transferred to nitrocellulose membranes and blocked overnight at 4°C or for 1 hour at room temperature in 5% non-fat dry milk/TBS. Antibodies were diluted in 5% BSA/TBS as follows: rabbit anti-Cx43 (Cat. No. 3512; Cell Signaling) at 1:2000 and mouse anti-Vinculin (Cat. No. V9131; Sigma) at 1:5000. Blots were incubated with primary antibodies overnight at 4°C or for 2 hours at room temperature then washed with TBST. Secondary HRP-conjugated anti-rabbit or anti-mouse antibodies were diluted at 1:5000 in TBST and incubated overnight at 4°C or for 2 hours at room temperature. Blots were incubated in ECL buffer (100 mM Tris, pH 8.8, 2.5 mM luminol, 0.4 mM p-coumaric acid, 0.02 % hydrogen peroxide) prior to exposure to film. ImageJ software was used to measure relative densities normalized to vinculin. Cx43 protein intensities were quantified with ImageJ using the gel analyzer tool and normalized to corresponding vinculin intensities for three separately transfected cultures expressing each transfected construct

### Scrape loading dye transfer assay

MDCK cells were seeded into 3.5 cm dishes and grown to ∼70% confluency. Dishes were transfected with WT Cx43, Cx43^Δ256-289^ constructs as described above. One set of dishes were left as untransfected controls. Transfection efficiency was estimated by transfecting a Cx43-GFP expressing construct (as described in Falk, 2000) concurrently. All transfection efficiencies were 80% or higher. At ∼24 hours post transfection, cells (∼100% confluent) were washed once with 1X PBS and 1mL of 0.05 weight % Lucifer Yellow (Cat. No L682; Invitrogen) in 1X PBS was added to the culture medium of each dish. Monolayers were scraped with a sharp razor blade and incubated at 37°C, under 5% CO_2_, and 100% humidity for 10 minutes. Cells were rinsed with 1X PBS and fixed in 3.7% formaldehyde for 10 minutes at room temperature. Cells were rinsed two times in 1X PBS and imaged with a 10X objective on a Nikon Eclipse TE2000 inverted fluorescence microscope. Images were analyzed by measuring the distance the dye transferred away from the scrape (10 measurements/scrape, 3 scrapes/ transfected dish, 3 separate transfected cultures expressing each construct) in ImageJ.

### RNA extraction and qRT-PCR analyses

Total RNA was extracted and purified from euthanized, unfixed, 3-month-old zebrafish (1 fish/sample) using the RNeasy Mini Kit (Cat. No. 74104; Qiagen) per the manufacturer’s instructions. Fish were homogenized by bead beating in the RLT buffer (RNeasy Kit) supplemented with 10µL BME/mL. qRT-PCR was performed using the QuantiNova SYBR Green RT-PCR kit (Cat. No. 208154; Qiagen) and CT values were measured using the Rotor-Gene (Corbett) analyzer. The CT values for Cx43, (forward: TCGCGTACTTGGATTTGGTGA, reverse: CCTTGTCAAGAAGCCTTCCCA) VEGF, (forward: TGCTCCTGCAAATTCACACAA, reverse: ATCTTGGCTTTTCACATCTGCAA) and the internal control, keratin (forward: TCATCGACAAAGTGCGCTTC, reverse: TCGATGTTGGAACGTGTGGT), were averaged. The ΔC_T_ values represent the expression levels of the gene of interest normalized to the internal control, keratin. ΔΔC_T_ values represent the level of gene expression. The fold difference was determined using the ΔΔC_T_ method as previously described (Livak *et al*., 2001; 2008). Three technical replicates/gene were performed for each of the 5 biological replicates.

### Heart rate analyses

At 3 and 6dpf, zebrafish embryos were collected for heart rate analyses. Individual embryos were placed on a glass coverslip in a drop of water. Heart rates were manually determined by counting individual beats visualized under a light microscope for a total of 15 seconds each, using a cell counter. This was performed 3 times for each embryo (100 for each genotype). Beats/15s were averaged for each embryo and multiplied in order to obtain the average beats/minute for each individual embryo. Images and movies of the heart were obtained with an iPhone X mounted to an Olympus inverted light microscope. Heart morphology was analyzed by tracing the atrium and ventricle of the heart on still frames from movies of the heartbeat using ImageJ.

### Vasculature analyses

Adult fin vasculature was analyzed in zebrafish that were anesthetized with 0.1% Tricaine solution and imaged using a Nikon Eclipse 80i Microscope (Diagnostic Instruments). Movies of fish were taken using a Levenhuk M300 BASE Camera and Levenhuk software. In addition to movies, three images/fin were obtained, and three diameter measurements were acquired/vessel for each image in ImageJ. In addition, blood flow distance/second was measured in ImageJ by tracing the traveled path of a single red blood cell through the vasculature for one second and measuring the distance traveled. One measurement was taken/vessel for each movie. Juvenile vasculature was analyzed by crossing *cx43^lh10^* with the *Tg*(*fli1:egfp*) transgenic line that drives GFP expression in all vasculature throughout development (Lawson et al., 2002) so the resultant progeny are *cx43^lh10^* zebrafish with GFP expressing vasculature. *Tg*(*fli1:egfp*)/*cx43^lh10^* zebrafish were fixed in 4% formaldehyde at 5dpf, and z-stacks were generated with confocal microscopy. Vasculature diameter for all intersegmental vessels were measured using ImageJ in five (WT) or three (*cx43^lh10^*) biological replicates.

### Statistical analyses

Statistical significance was determined by student’s T-test (two-tailed, unpaired). Statistical significance was defined by a p-value of ≤ 0.05.

## ACKNOWLEDGEMENTS

We thank current and previous Falk and Iovine lab members for constructive discussions. This work was supported by the National Institute of Health’s Institute of General Medical Sciences Grant GM55725 to M.M.F and Lehigh University’s Faculty Innovation Grant.

## Abbreviations

Cx: Connexin
Cx43: Connexin43
GJIC: gap junction intercellular communication
CT: C-terminal
WT: wild type

---

MOVIE 1: Normal cardiac beat cycle in WT zebrafish at 3dpf.

MOVIE 2: Bradycardia in *cx43^lh10^* zebrafish at 3dpf also exhibiting pericardial edema.

MOVIE 3: Normal vasculature organization and blood flow in WT adult zebrafish caudal fins. Distal (top) and Proximal (bottom). Arteries are located in the center of the fin ray bone, while veins flank the bone on either side.

MOVIE 4: Abnormal vasculature organization in *cx43^lh10^* adult zebrafish caudal fins. Distal (top) and Proximal (bottom). In addition to disorganized vasculature, it is also apparent that some vessels have dramatically reduced blood flow.

MOVIE 5: Dramatically reduced blood flow in *cx43^lh10^* adult zebrafish caudal fins. Distal (top) and Proximal (bottom).

## Notes

### Competing Interest Statement

The authors have declared no competing interest.

## REFERENCES

1. Bakkers J (2011). Zebrafish as a Model to Study Cardiac Development and Human Cardiac Disease. Cardiovasc Res 91, 279–288.

2. Beardslee MA, Laing JG, Beyer EC, Saffitz JE (1998). Rapid Turnover of connexin43 in the Adult Rat Heart. Circ Res 83, 629–635.

3. Beardslee MA, Lerner DL, Tadros PN, Laing JG, Beyer EC, Yamada KA, Kléber AG, Schuessler RB, Saffitz JE (2000). Dephosphorylation and Intracellular Redistribution of Ventricular Connexin43 during Electrical Uncoupling Induced by Ischemia. Circ Res 87, 656–662.

4. Bedner P, Steinhäuser C, Theis M. (2012). Functional Redundancy and Compensation among Members of Gap Junction Protein Families? Biochim Biophys Acta Biomembr 1818, 1971–1984.

5. Bhattacharya S, Hyland C, Falk, MM, Iovine MK (2020). Connexin 43 Gap Junctional Intercellular Communication Inhibits Evx1 Expression and Joint Formation in Regenerating Fins. Development 147.

6. Brehm R, Zeiler M, Rüttinger C, Herde K, Kibschull M, Winterhager E, Willecke K, Guillou F, Lécureuil C, Steger K et al. (2007). A Sertoli Cell-Specific Knockout of connexin43 Prevents Initiation of Spermatogenesis. Am J Pathol 171, 19–31.

7. Bruce AF, Rothery S, Dupont E, Severs NJ (2008). Gap Junction Remodelling in Human Heart Failure is Associated with Increased Interaction of connexin43 with ZO-1. Cardiovasc Res 77, 757–765.

8. Chatterjee B, Chin AJ, Valdimarsson G, Finis C, Sonntag JM, Choi BY, Tao L, Balasubramanian K, Bell C, Krufka A et al. (2005). Developmental Regulation and Expression of the Zebrafish connexin43 Gene. Dev Dyn 233, 890–906.

9. Chi NC, Bussen M, Brand-Arzamendi K, Ding C, Olgin JE, Shaw RM, Martin GR, Stainier DYR (2010). Cardiac Conduction is Required to Preserve Cardiac Chamber Morphology. Proc Natl Acad Sci U S A 107, 14662–14667.

10. Chu FF, Doyle D (1985) Turnover of plasma membrane proteins in rat hepatoma cells and primary cultures of rat hepatocytes. J Biol Chem 260: 3097–3107.

11. Cofre J, Abdelhay E (2007). Connexins in the Early Development of the African Clawed Frog Xenopus Laevis (Amphibia): The Role of the connexin43 Carboxyl Terminal Tail in the Establishment of the Dorso-Ventral Axis. Genet Mole Biol 30, 483–493.

12. Cone AC, Cavin G, Ambrosi C, Hakozaki H, Wu-Zhang AX, Kunkel MT, Newton AC, Sosinsky GE (2014). Protein Kinase Cδ-Mediated Phosphorylation of Connexin43 Gap Junction Channels Causes Movement within Gap Junctions Followed by Vesicle Internalization and Protein Degradation. J Biol Chem 289, 8781–8798.

13. Duffy HS (2012). The Molecular Mechanisms of Gap Junction Remodeling. Heart Rhythm 9, 1331–1334.

14. Dunn CA, Lampe PD (2014). Injury-Triggered Akt Phosphorylation of Cx43: A ZO-1-Driven Molecular Switch that Regulates Gap Junction Size. J Cell Sci 127, 455–464.

15. Falk MM (2000). Connexin-Specific Distribution within Gap Junctions Revealed in Living Cells. J Cell Sci 113, 4109–4120.

16. Falk MM, Baker SM, Gumpert AM, Segretain D, Buckheit RW (2009). Gap Junction Turnover is Achieved by the Internalization of Small Endocytic Double-Membrane Vesicles. Mol Biol Cell 20, 3342–3352.

17. Falk MM, Kells RM, Berthoud VM (2014). Degradation of Connexins and Gap Junctions. FEBS Lett 588, 1221–1229.

18. Falk MM, Fong J, Kells R, O’Laughlin M, Kowal T, Thévenin A (2012). Degradation of Endocytosed Gap Junctions by Autophagosomal and Endo-/Lysosomal Pathways: A Perspective. J Membr Biol 245, 465–476.

19. Fallon RF, Goodenough DA (1981). Five-Hour Half-Life of Mouse Liver Gap-Junction Protein. J Cell Biol 90, 521–526.

20. Figueroa XF, Duling BR (2008). Gap Junctions in the Control of Vascular Function. Antioxid Redox Signal 11, 251–266.

21. Figueroa XF, Isakson BE, Duling BR (2006). Vascular Gap Junctions in Hypertension. Hypertension 48, 804–811.

22. Fong JT, Kells RM, Falk MM (2013). Two Tyrosine-Based Sorting Signals in the Cx43 C-Terminus Cooperate to Mediate Gap Junction Endocytosis. Mol Biol Cell 24, 2834–2848.

23. Fong JT, Kells RM, Gumpert AM, Marzillier JY, Davidson MW, Falk MM (2012). Internalized Gap Junctions are Degraded by Autophagy. Autophagy 8, 794–811.

24. Fong JT, Nimlamool W, Falk MM (2014). EGF Induces Efficient Cx43 Gap Junction Endocytosis in Mouse Embryonic Stem Cell Colonies Via Phosphorylation of Ser262, Ser279/282, and Ser368. FEBS Lett 588, 836–844.

25. Girão H, Catarino S, Pereira P (2009). Eps15 Interacts with Ubiquitinated Cx43 and Mediates its Internalization. Exp Cell Res 315, 3587–3597.

26. Gumpert AM, Varco JS, Baker SM, Piehl M, Falk MM (2008). Double-Membrane Gap Junction Internalization Requires the Clathrin-Mediated Endocytic Machinery. FEBS Lett 582, 2887–2892.

27. Gutstein DE, Morley GE, Tamaddon H, Vaidya D, Schneider MD, Chen J, Chien KR, Stuhlmann H, Fishman GI (2001). Conduction Slowing and Sudden Arrhythmic Death in Mice with Cardiac-Restricted Inactivation of connexin43. Circ Res 88, 333–339.

28. Haefliger J, Nicod P, Meda P (2004). Contribution of Connexins to the Function of the Vascular Wall. Cardiovasc Res 62, 345–356.

29. Hardy K, Warner A, Winston RML, Becker DL (1996). Expression of Intercellular Junctions during Preimplantation Development of the Human Embryo. Mol Hum Reprod 2, 621–632.

30. Hare JF and Taylor K (1991). Mechanisms of plasma membrane protein degradation: recycling proteins are degraded more rapidly than those confined to the cell surface. Proc Natl Acad Sci U S A 88: 5902–5906,

31. Inoguchi T, Yu HY, Imamura M, Kakimoto M, Kuroki T, Maruyama T, Nawata H (2001). Altered Gap Junction Activity in Cardiovascular Tissues of Diabetes. Med Electron Microsc 34, 86–91.

32. Karczewski KJ, Francioli LC, Tiao G, Cummings BB, Alföldi J, Wang Q, Collins RL, Laricchia KM, Ganna A, Birnbaum DP et al. (2020). The Mutational Constraint Spectrum Quantified from Variation in 141,456 Humans. Nature 581, 434–443.

33. Kells-Andrews RM, Margraf RA, Fisher CG, Falk MM (2018). Connexin-43 K63-Polyubiquitylation on Lysines 264 and 303 Regulates Gap Junction Internalization. J Cell Sci 131.

34. Kwak BR, Pepper MS, Gros DB, Meda P (2001). Inhibition of Endothelial Wound Repair by Dominant Negative Connexin Inhibitors. Mol Biol Cell 12, 831–845.

35. Labun K, Montague TG, Gagnon JA, Thyme SB, Valen E (2016). CHOPCHOP V2: A Web Tool for the Next Generation of CRISPR Genome Engineering. Nucleic Acids Res 44, W272–W276.

36. Labun K, Montague TG, Krause M, Torres Cleuren YN, Tjeldnes H, Valen E (2019). CHOPCHOP V3: Expanding the CRISPR Web Toolbox Beyond Genome Editing. Nucleic Acids Res 47, W171–W174.

37. Lampe PD, Lau AF (2004). The Effects of Connexin Phosphorylation on Gap Junctional Communication. Int J Biochem Cell Biol 36, 1171–1186.

38. Lawson ND, Weinstein BM (2002). In vivo imaging of embryonic vascular development using transgenic zebrafish. Dev. Biol. 248, 307–318.

39. Leithe E, Mesnil M, Aasen T (2018). The Connexin 43 C-Terminus: A Tail of Many Tales. Biochim Biophys Acta Biomembr 1860, 48–64.

40. Lek M, Karczewski KJ, Minikel EV, Samocha KE, Banks E, Fennell T, O’Donnell-Luria AH, Ware JS, Hill AJ, Cummings BB, et al. (2016). Exome Aggregation C. Analysis of protein-coding genetic variation in 60,706 humans. Nature 536: 285–291,

41. Livak KJ, Schmittgen TD (2001). Analysis of Relative Gene Expression Data using Real-Time Quantitative PCR and the 2(-Delta Delta C(T)) Method. Methods 25, 402–408.

42. Livak KJ, Schmittgen TD (2008). Analyzing Real-Time PCR Data by the Comparative C T Method. Nat Protoc 3, 1101-1108.

43. Lübkemeier I, Requardt R, Lin X, Sasse P, Andrié R, Schrickel J, Chkourko H, Bukauskas F, Kim J, Frank M et al. (2013). Deletion of the Last Five C-Terminal Amino Acid Residues of connexin43 Leads to Lethal Ventricular Arrhythmias in Mice without Affecting Coupling Via Gap Junction Channels. Basic Res Cardiol 108, 1–16.

44. Maass K, Chase SE, Lin X, Delmar M (2009). Cx43 CT Domain Influences Infarct Size and Susceptibility to Ventricular Tachyarrhythmias in Acute Myocardial Infarction. Cardiovasc Res 84, 361–367.

45. Maass K, Shibayama J, Chase SE, Willecke K, Delmar M (2007). C-Terminal Truncation of Connexin43 Changes Number, Size, and Localization of Cardiac Gap Junction Plaques. Circ Res 101, 1283–1291.

46. Martins-Marques T, Anjo SI, Pereira P, Manadas B, Girão H (2015a). Interacting Network of the Gap Junction (GJ) Protein Connexin43 (Cx43) is Modulated by Ischemia and Reperfusion in the Heart. Mol Cell Proteomics 14, 3040–3055.

47. Martins-Marques T, Catarino S, Marques C, Matafome P, Ribeiro-Rodrigues T, Baptista R, Pereira P, Girão H (2015b). Heart Ischemia Results in connexin43 Ubiquitination Localized at the Intercalated Discs. Biochimie 112, 196–201.

48. Martins-Marques T, Catarino S, Marques C, Pereira P, Girão H (2015c). To Beat or Not to Beat: Degradation of Cx43 Imposes the Heart Rhythm. Biochem Soc Trans 43, 476–481.

49. Martins-Marques T, Catarino S, Zuzarte M, Marques C, Matafome P, Pereira P, Girão H (2015d). Ischaemia-Induced Autophagy Leads to Degradation of Gap Junction Protein connexin43 in Cardiomyocytes. Biochem J 467, 231–245.

50. McLachlan E, Shao Q, Wang H, Langlois S, Laird DW (2006). Connexins Act as Tumor Suppressors in Three-Dimensional Mammary Cell Organoids by Regulating Differentiation and Angiogenesis. Cancer Res 66, 9886–9894.

51. Michela P, Velia V, Aldo P, Ada P (2015). Role of Connexin 43 in Cardiovascular Diseases. Eur J Pharmacol 768, 71–76.

52. Montague TG, Cruz JM, Gagnon JA, Church GM, Valen E (2014). CHOPCHOP: A CRISPR/Cas9 and TALEN Web Tool for Genome Editing. Nucleic Acids Res 42, W401–W407.

53. Nimlamool W, Andrews RMK, Falk MM (2015). Connexin43 Phosphorylation by PKC and MAPK Signals VEGF-Mediated Gap Junction Internalization. Mol Biol Cell 26, 2755–2768.

54. Park DJ, Freitas TA, Wallick CJ, Guyette CV, Warn-Cramer BJ (2006). Molecular Dynamics and in Vitro Analysis of Connexin43: A New 14-3-3 Mode-1 Interacting Protein. Protein Sci 15, 2344–2355.

55. Park DJ, Wallick CJ, Martyn KD, Lau AF, Jin C, Warn-Cramer BJ (2009). Akt Phosphorylates Connexin43 on Ser373, a “Mode-1” Binding Site for 14-3-3. Cell Commun Adhes 14, 211–226.

56. Pepper MS, Meda P (1992). Basic Fibroblast Growth Factor Increases Junctional Communication and Connexin 43 Expression in Microvascular Endothelial Cells. J Cell Physiol 153, 196–205.

57. Pepper MS, Montesano R, el Aoumari A, Gros D, Orci L, Meda P (1992). Coupling and Connexin 43 Expression in Microvascular and Large Vessel Endothelial Cells. Am J Physiol 262, 1246-1257.

58. Piehl M, Lehmann C, Gumpert A, Denizot J, Segretain D, Falk MM (2007). Internalization of Large Double-Membrane Intercellular Vesicles by a Clathrin-Dependent Endocytic Process. Mol Biol Cell 18, 337–347.

59. Plum A, Hallas G, Magin T, Dombrowski F, Hagendorff A, Schumacher B, Wolpert C, Kim J, Lamers WH, Evert M et al. (2000). Unique and Shared Functions of Different Connexins in Mice. Curr Biol 10, 1083–1091.

60. Reaume AG, de Sousa PA, Kulkarni S, Langille BL, Zhu D, Davies TC, Juneja SC, Kidder GM, Rossant J (1995). Cardiac Malformation in Neonatal Mice Lacking connexin43. Science 267, 1831–1834.

61. Rhett JM, Jourdan J, Gourdie RG (2011). Connexin 43 Connexon to Gap Junction Transition is Regulated by Zonula Occludens-1. Mol Biol Cell 22, 1516–1528.

62. Rummery NM, Hill CE (2004). Vascular Gap Junctions and Implications for Hypertension. Clin Exp Pharmacol Physiol 31, 659–667.

63. Sridharan S, Simon L, Meling DD, Cyr DG, Gutstein DE, Fishman GI, Guillou F, Cooke PS. (2007). Proliferation of Adult Sertoli Cells Following Conditional Knockout of the Gap Junctional Protein GJA1 (Connexin 43) in Mice. Biol Reprod 76, 804–812.

64. Schindelin J, Arganda-Carreras I, Frise E, Kaynig V, Longair M, Pietzsch T, Preibisch S, Rueden C, Saalfeld S, Schmid B et al. (2012). Fiji: An Open-Source Platform for Biological-Image Analysis. Nat Methods 9, 676–682.

65. Severs NJ, Coppen SR, Dupont E, Yeh H, Ko Y, Matsushita T (2004). Gap Junction Alterations in Human Cardiac Disease. Cardiovasc Res 62, 368–377.

66. Singleman C, Holtzman NG (2011). Heart Dissection in Larval, Juvenile and Adult Zebrafish, Danio Rerio. J Vis Exp 55, 3165.

67. Solan JL, Lampe PD (2007). Key Connexin 43 Phosphorylation Events Regulate the Gap Junction Life Cycle. J Membr Biol 217, 35–41.

68. Suarez S, Ballmer-Hofer K (2001). VEGF Transiently Disrupts Gap Junctional Communication in Endothelial Cells. J Cell Sci 114, 1229–1235.

69. Thévenin AF, Kowal TJ, Fong JT, Kells RM, Fisher CG, Falk MM (2013). Proteins and Mechanisms Regulating Gap-Junction Assembly, Internalization, and Degradation. Physiology (Bethesda) 28, 93–116.

70. Thévenin AF, Margraf RA, Fisher CG, Kells-Andrews RM, Falk MM (2017). Phosphorylation Regulates connexin43/ZO-1 Binding and Release, an Important Step in Gap Junction Turnover. Mol Biol Cell 28, 3595–3608.

71. Wang W, Chen M, Leong H, Kuo Y, Kuo C, Lee C (2014). Connexin 43 Suppresses Tumor Angiogenesis by Down-Regulation of Vascular Endothelial Growth Factor Via Hypoxic-Induced Factor-1α. Int J Mol Sci 16, 439–451.

72. Winterhager E, Pielensticker N, Freyer J, Ghanem A, Schrickel JW, Kim J, Behr R, Grümmer R, Maass K, Urschel S et al. (2007). Replacement of connexin43 by connexin26 in Transgenic Mice Leads to Dysfunctional Reproductive Organs and Slowed Ventricular Conduction in the Heart. BMC Dev Biol 7, 26.

73. Ya J, Erdtsieck-Ernste EB, de Boer PA, van Kempen MJ, Jongsma H, Gros D, Moorman AF, Lamers WH (1998). Heart Defects in connexin43-Deficient Mice. Circ Res 82, 360–366.

74. Zhang J, Hill CE (2005). Differential Connexin Expression in Preglomerular and Postglomerular Vasculature: Accentuation during Diabetes. Kidney Int 68, 1171–1185.

