## Supplemental Figures for "Impaired Cx43 gap junction endocytosis causes cardiovascular defects in zebrafish"

Supplemental Table 1: Quantitative RT-PCR results. The ∆C_T_ value is determined by subtracting the average C_T_ value of a housekeeping gene (keratin) from the average gene C_T_ value. The standard deviation of the difference is calculated from the standard deviations of the gene and keratin values using the comparative method. The calculation of the ∆∆C_T_ involves subtraction of the ∆C_T_ calibrator value. This is a subtraction of an arbitrary constant, so the deviation of the ∆∆C_T_ is the same as the standard deviation of the ∆C_T_ value. The range given for a gene relative to WT is determined by evaluating the expression 2^-∆∆CT^ with ∆∆C_T_ -s, where s= standard deviation of the ∆∆CT value. At least three biological replicates were performed for all comparisons.

Supplemental Figure 1: Verification of the *cx43^lh10^* deletion and generation of the *cx43^lh10^* zebrafish line. (A) Representative agarose gel showing the PCR verification of CRISPR/Cas9 injected zebrafish embryos (24hpf). WT allele is represented by a ~550bp band, whereas the *cx43^lh10^* allele is represented by a ~450bp band. (B) Representative agarose gel showing the PCR verification of a heterozygous intercross. (C) Homozygous *cx43^lh10^* zebrafish line generation from injection to heterozygous cross breading. One-cell stage embryos were injected with CRISPR components, generating mosaic embryos (not all cells of the embryo contain a copy of the mutant allele), as represented by the gel in (A). Mosaic adults were outcrossed to WT zebrafish for germline transmission and generation of heterozygous fish (all cells contain one copy of the mutant allele). Heterozygous adults were intercrossed for generation of homozygous *cx43^lh10^* zebrafish (all cells contain 2 copies of the mutant allele), as represented by the gel in (B).

Supplemental Figure 2: *cx43^lh10^* zebrafish embryos show no obvious morphological changes or altered viability during early stages of development. (A) Early developmental morphology of WT (top) and *cx43^lh10^* embryos (bottom). (B) Viability of WT and *cx43^lh10^* zebrafish over time. (n=270)

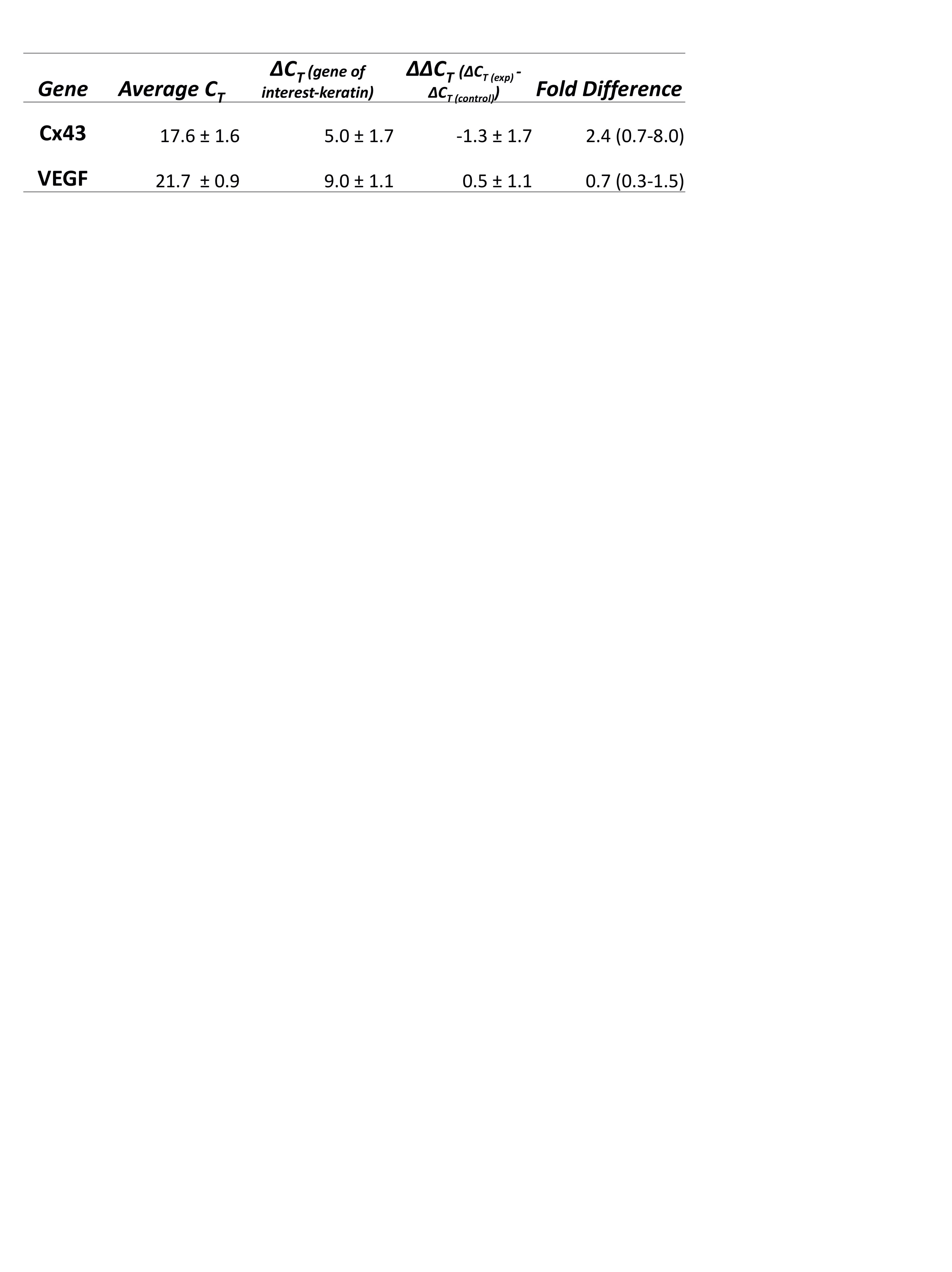

Supplemental Table 1

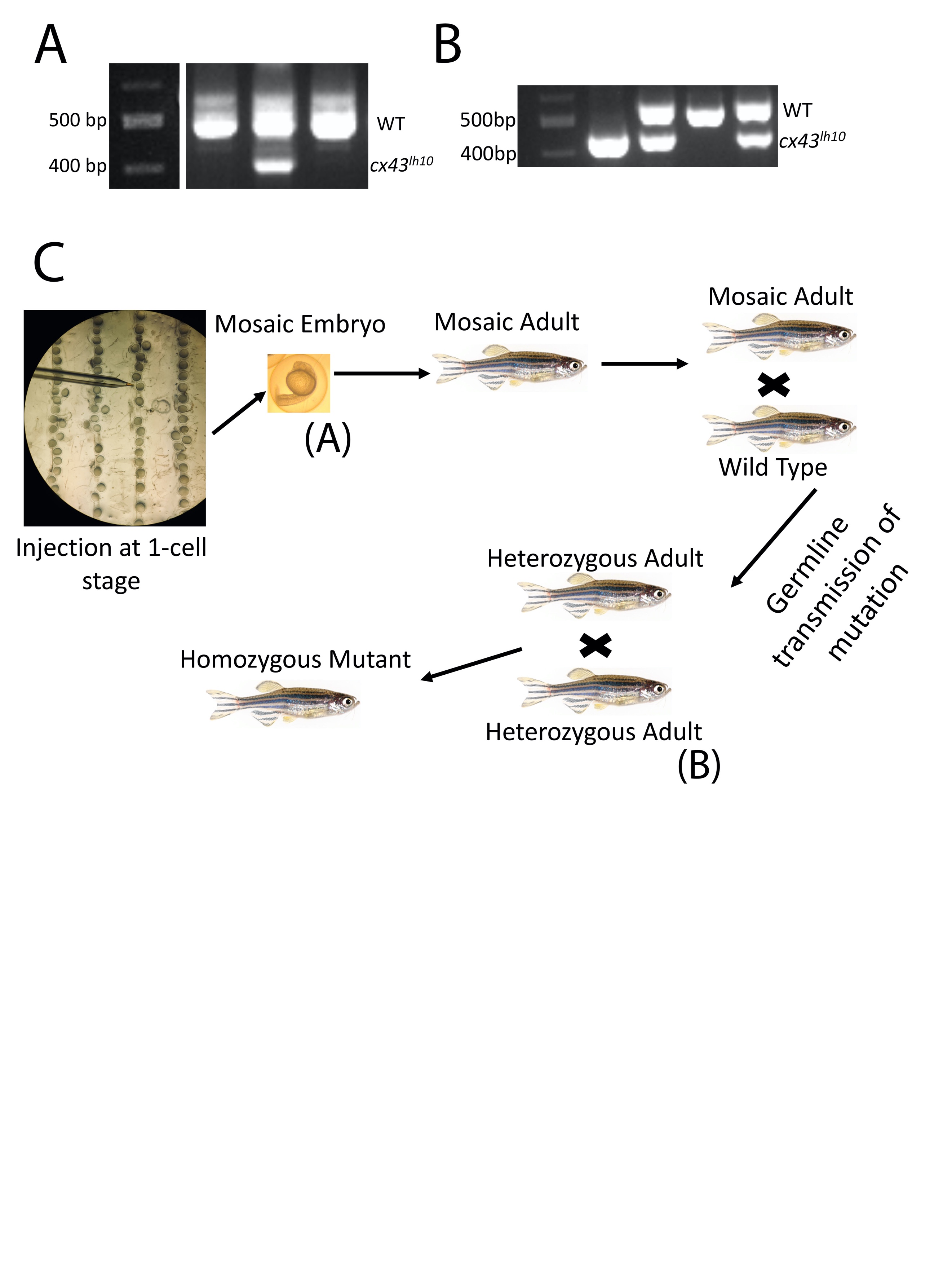

Supplemental Fig 1

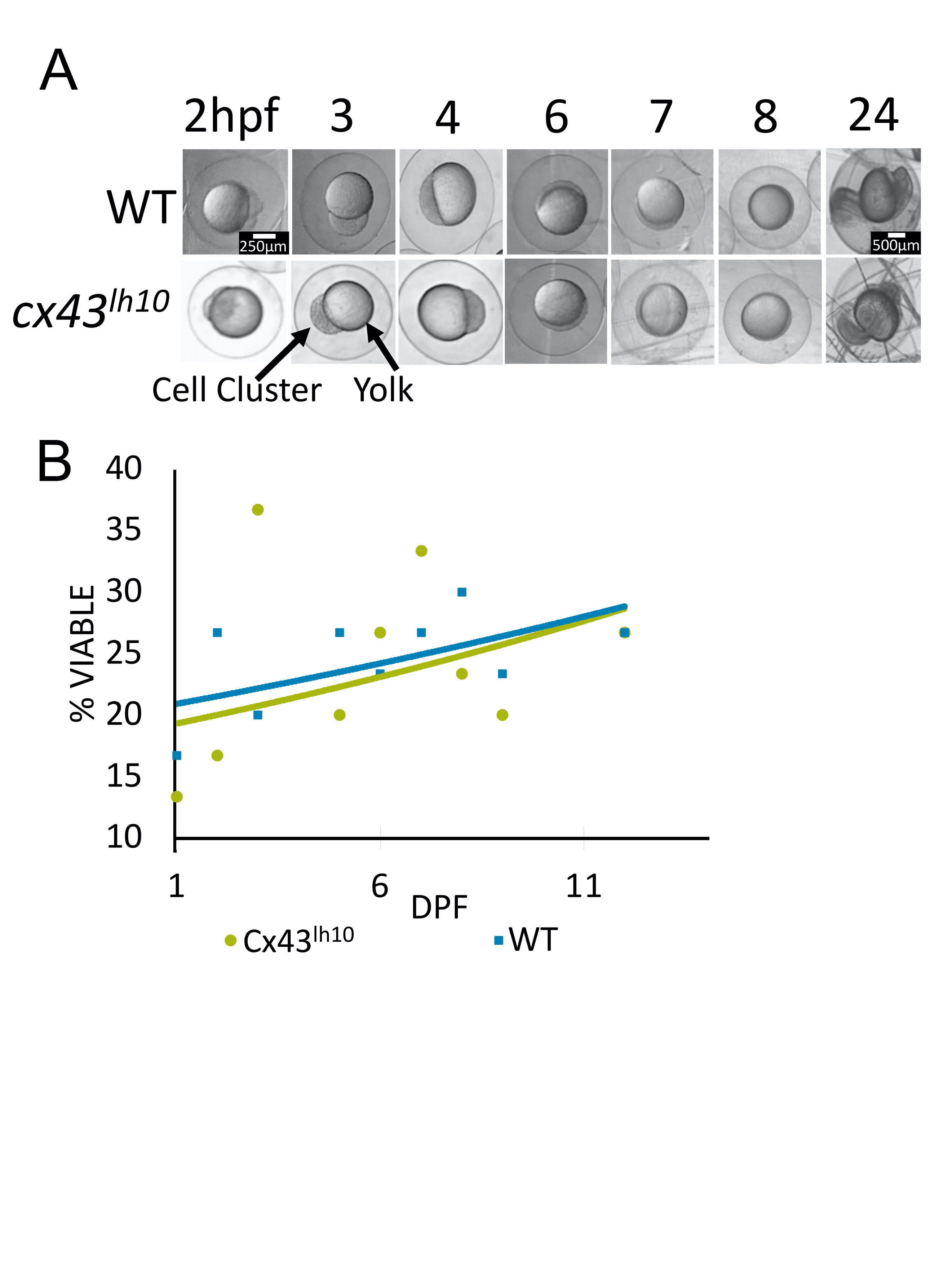

Supplemental Figure 2
